# FIRST-seq: a nanopore-based cDNA sequencing platform for RNA modification and structure profiling

**DOI:** 10.1101/2025.04.22.649934

**Authors:** Oguzhan Begik, Gregor Diensthuber, Ivana Borovska, John S Mattick, Danny Incarnato, Eva Maria Novoa

**Author notes:** Correspondence to: Eva Maria Novoa.

## Abstract

RNA modifications induce reverse transcription (RT) errors in an enzyme- and context-dependent manner, enabling transcriptome-wide mapping and RNA structure probing. We present FIRST-seq, a flexible, cost-effective nanopore cDNA method that avoids second-strand synthesis and PCR, making it compatible with any RT enzyme and enabling single-nucleotide resolution RT signature analysis. Benchmarking multiple RT enzymes and buffers identified conditions that reduce premature termination and enhance error detection. Coupled with DMS probing, FIRST-seq accurately detects m1A and m3C at unpaired sites, recapitulating known RNA structures in vitro and in vivo. FIRST-seq offers a versatile platform for profiling chemical-induced and natural RNA modifications using long-read sequencing.

## INTRODUCTION

In recent years, highly-processive Group II intron reverse transcriptases have been increasingly adopted in transcriptomic research, due to their ability to generate full-length, complementary DNA (cDNA) fragments from RNA templates with high fidelity, even in the presence of extensive secondary structures or post-transcriptional modifications [1–9]. The development of numerous library preparation protocols incorporating these enzymes has established them as essential tools in transcriptome-wide studies utilizing next-generation sequencing (NGS) technologies. Their widespread application includes: i) generation of comprehensive long and short RNA-seq libraries, as exemplified by protocols such as TGIRT-seq [4,10], mim-tRNA-seq [2] and OTTR-seq [11]; ii) the detection of endogenous RNA modifications, typically manifested as increased errors and mismatch rates during reverse transcription [3,12–16]; and iii) the generation of transcriptome-wide RNA structural profiles, obtained through chemical probing coupled with NGS, as implemented in techniques such as DMS-MaP-Seq [17] or SHAPE-MaP [18].

With ongoing improvements in the accuracy of third generation sequencing technologies (TGS) –mainly represented by Pacific Biosciences (PacBio) [19] and Oxford Nanopore Technologies (ONT) [20–22]–, several methodologies originally designed for next-generation sequencing are increasingly being adapted for the use with TGS platforms [23–26]. Despite this progress, the integration of highly-processive RT enzymes to TGS-based workflows remains limited, primarily due to their incompatibility with currently available commercial library preparation kits.

To overcome this limitation, here we develop FIRst STrand sequencing (FIRST-seq), a novel nanopore-based cDNA sequencing method that enables direct sequencing of the first-strand synthesized cDNA without the need of second-strand synthesis or PCR amplification, thus being compatible with any RT enzyme of interest, without the need of additional protocol adaptations. We first systematically evaluate the performance of FIRST-seq with a range of RT enzymes under varying buffer conditions, identifying specific combinations that yield highest RT mismatch error signatures while minimizing the reverse transcription drop-off. We first demonstrate its applicability at identifying known endogenous m^1^A modifications in human RNAs. We then couple dimethyl sulphate (DMS) –a chemical probing reagent commonly used for RNA structure profiling [17,22,27–29]– with FIRST-seq, demonstrating its ability to generate high-accuracy, transcriptome-wide RNA structure maps both *in vivo* and *in vitro*, overcoming the need for gene-specific primers [22]. Notably, FIRST-seq can be coupled with any chemical probing agent of interest that induces mismatch errors or RT drop-offs, thereby facilitating the adaptation of existing NGS-based methods that employ highly-processive RT enzymes (e.g., TGIRT, Induro RT, Marathon RT) for use with nanopore sequencing platforms. Overall, FIRST-seq expands the applicability of nanopore sequencing by enabling the detection of natural and chemically-induced RNA modifications using cDNA-based methodologies.

## RESULTS

### Modified nucleotides cause wide signal shifts in direct RNA sequencing data, but not cDNA

In contrast to NGS methods, the nanopore sequencing platform is capable of sequencing not only cDNA but also native RNA molecules directly, thereby eliminating the need for reverse transcription [30–36]. This capacity enables coupling of chemical probing with direct RNA sequencing (dRNA-seq) in addition to the conventional approach of coupling with cDNA-based sequencing (**Figure 1a**). In this regard, previous works have demonstrated that dRNA-seq can successfully capture RNA structural information when coupled with chemical probing [20]. However, we hypothesized that this approach might be suboptimal for detecting chemical-probing signals, as RNA modifications are known to cause perturbations in current intensity and basecalling ‘errors’ not only at the modified site but also at adjacent nucleotides [21,37–39], thereby increasing background noise and reducing signal resolution for RNA structure predictions (**Figure 1b**).

**Figure 1.**
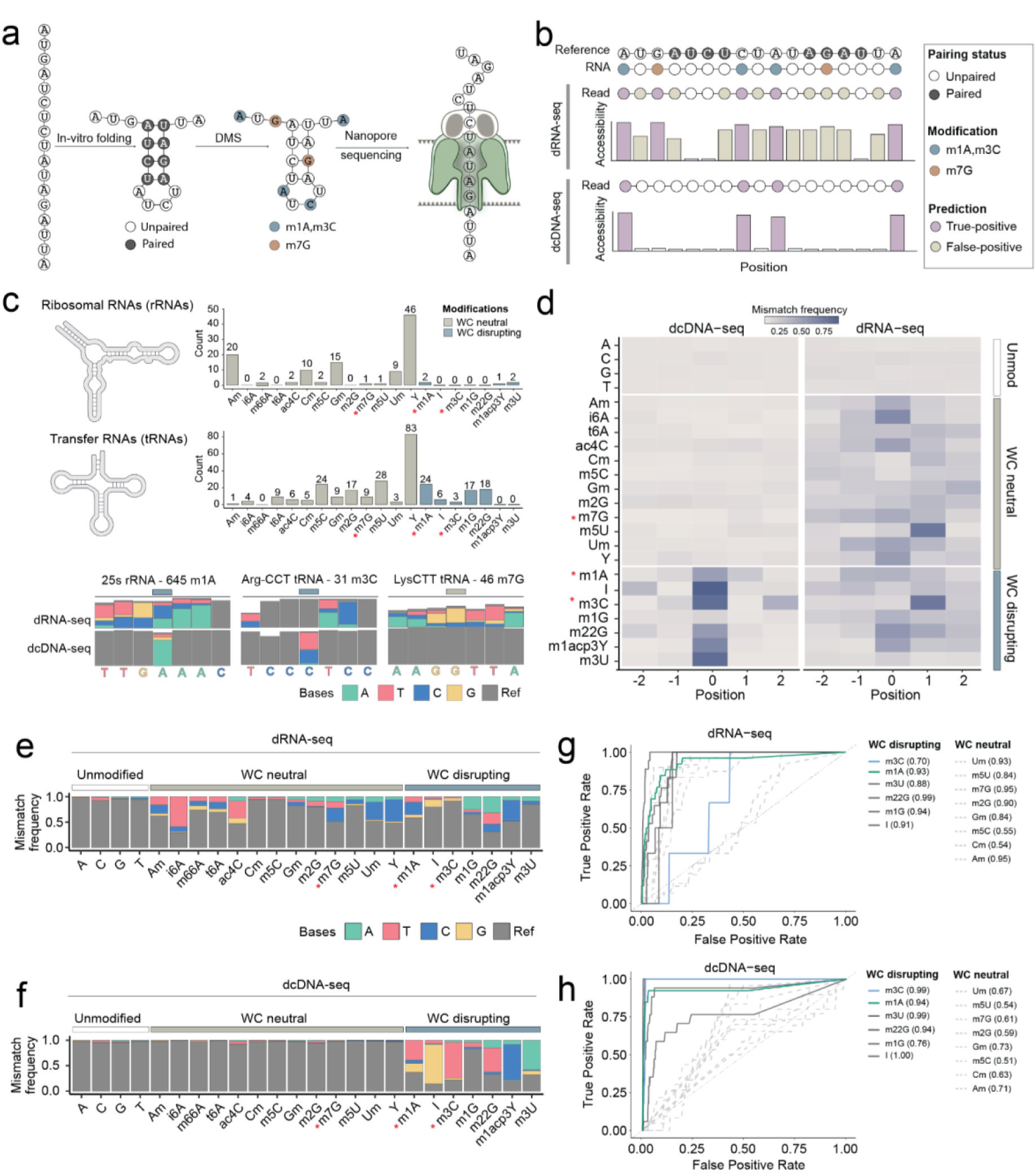
Most RNA modifications lead to single nucleotide error signatures in direct cDNA sequencing, whereas they produce ‘spread’ error signals in direct RNA sequencing datasets. **(a)** Schematic overview of the DMS probing coupled to nanopore sequencing, which can be sequenced either in its native RNA form (dRNA-seq) or after reverse transcription (dcDNA-seq). Gray-colored circles indicate paired bases, while white circles represent unpaired bases. Dimethyl sulfate (DMS) treatment methylates unpaired A and C positions, as well as both paired and unpaired G positions. Methylation leads to the formation of m^1^A and m^3^C modifications (blue) and m^7^G modifications (orange). **(b)** Schematic representation of how DMS-induced modifications affect the signal in the RNA molecule and consequently the mapped accessibility. While dRNA-seq detects modified sites (shown in purple), the “error” signature often spans to neighboring unmodified bases (shown in brown), resulting in a higher false positive rate. In contrast, direct cDNA sequencing (dcDNA-seq) produces “error” signatures only from WC-disrupting modifications (m^1^A, m^3^C), ignoring m^7^G, which is modified regardless of its pairing status. Consequently, cDNA reads from DMS-probed RNAs are expected to have a higher true positive error rate. **(c)** Ribosomal RNAs (rRNAs) and transfer RNAs (tRNAs) are heavily modified (top panel). In the bottom panel, Integrated Genomics Viewer (IGV) coverage track snapshots exemplify dRNA-seq and dcDNA-seq data at three modified positions Colored bars on top of the barplots indicate the modified positions. See also **Figure S1a** for additional IGV snapshots. **(d)** Heatmap showing aggregated basecalling error signatures along the 5-mer region (centered at the modified site, position 0) from multiple modified positions in tRNAs and rRNA molecules, sequenced with either dRNA-seq or dcDNA-seq. **See also Figure S1b,c**. **(e,f)** Barplot showing aggregated error signatures using dRNA-seq (e) and dcDNA-seq (f) data. DMS-induced modifications (m^1^A, m^3^C and m^7^G) are marked with a red star. **(g,h)** ROC curves depicting the true positive (TP) rate and false positive (FP) rate in detecting different RNA modifications using mismatch ‘error’ patterns, using either dRNA-seq (g) or dc-DNAseq (h). Watson-Crick (WC) disrupting bases are shown in full lines, with m^3^C and m^1^A highlighted in blue and green, respectively. WC-neutral bases are shown in dashed gray lines. The AUC value for each base and method is shown in parentheses in the legend. See also **Figure S2**.

Here we systematically evaluated whether native RNA (direct RNA sequencing; dRNA.seq) or cDNA-based (direct cDNA-sequencing; dcDNA-seq) nanopore sequencing offers greater accuracy in detecting diverse RNA modifications, including chemically-induced RNA modifications introduced by probing reagents, such as dimethyl sulphate (DMS). DMS selectively methylates unpaired adenosines (A) and cytosines (C), generating *N1*-methyladenosine (m^1^A), *N3*-methylcytosine (m^3^C) respectively. However, DMS also modifies both guanosines (G) at the N7 position to form N7-methylguanosine (m^7^G), regardless of their pairing status. This non-selective methylation can complicate RNA structure prediction by introducing noise and causing signal ‘leakage’ to neighboring positions [30,38], resulting in false positive predictions of structural accessibility (**Figure 1b**).

To assess whether cDNA-based detection provides higher resolution of chemically-induced modifications than native RNA sequencing, we analyzed the modification patterns in yeast rRNA and tRNAs, which contain well-characterized RNA modification landscapes (**Figure 1c**, see also **Figure S1a**), using both dRNA-seq and dcDNA-seq. Our results showed that Watson-Crick (WC)-disrupting RNA modifications, including m^1^A and m^3^C, resulted in single nucleotide resolution signatures in dcDNA-seq datasets, whereas WC-neutral modifications did not cause detectable changes in the base-calling patterns (**Figure 1d**, see also **Figure S1b**). Furthermore, RT misincorporation signatures observed in dcDNA-seq datasets were significantly stronger than ‘basecalling errors’ in dRNA-seq datasets –which can be used as a proxy for signal intensity differences [32–34,38]–at the corresponding modified sites (**Figure 1e-f**, see also **Figure S1b,c**). Collectively, these findings suggest that cDNA-based nanopore sequencing offers enhanced sensitivity and resolution for detecting specific chemically induced RNA modifications, such as m^3^C and m^1^A, compared to dRNA-seq (**Figure 1g-h**, see also **Table S1** and **S2**).

### FIRST-seq expands nanopore cDNA sequencing compatibility to all reverse transcriptases

The commercial direct cDNA sequencing (dcDNA-seq) and PCR-cDNA sequencing protocols offered by ONT (see *Methods*) are designed to capture full-length cDNA products by incorporating a strand-switching step, followed by second-strand synthesis [40]. The resulting double-stranded cDNA products (or their PCR-amplified equivalents) are then ligated to ONT-specific sequencing adapters (**Figure 2a**, left panel). While these methods are useful for RNA quantification and isoform detection [41,42], their use for RNA modification detection –particularly through RT mismatch or drop-off signatures– is limited. Indeed, these methods require the use of RT enzymes that add untemplated GGG overhangs at the 3’ end of the first-strand cDNA, used to initiate strand-switching and second-strand synthesis. While this enables efficient full-length cDNA capture, it limits the use of highly-processive RT enzymes, such as Group II intron RTs, which have been reported to outperform viral RTs in RNA modification detection and RNA structure probing applications due to their improved processivity and elevated misincorporation rates [8,17,22]. This limitation was recently addressed by employing gene-specific primers coupled with PCR amplification, allowing to couple the use of Marathon RT [9] –a highly processive group II intron RT enzyme– with nanopore cDNA sequencing, referred to as Nano-DMS-MaP-seq [22]. While effective, a major caveat of this strategy is that it is restricted to transcripts that are complementary to the gene-specific primers used at the reverse transcription step, and therefore is not suitable for transcriptome-wide RNA modification or RNA structure profiling.

**Figure 2.**
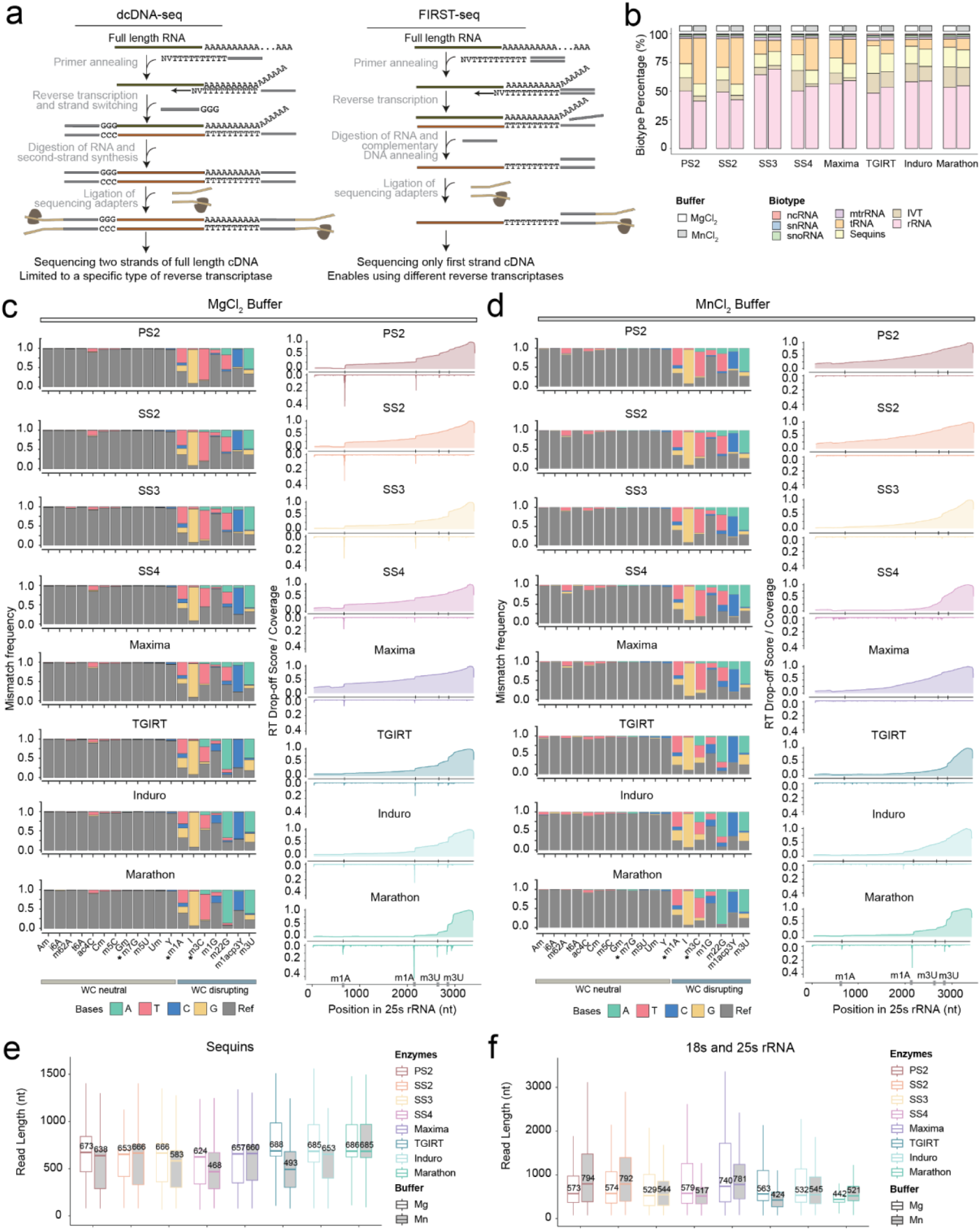
Comparison of reverse transcriptase (RT) performance in MgCl_2_ and MnCl_2_ buffers across different enzymes. **(a)** Schematic comparing workflows of dcDNA-seq and FIRST-seq. FIRST-seq simplifies the process by sequencing only the first cDNA strand, eliminating the need for strand-switching and second-strand synthesis. This allows for the use of a broader range of reverse transcriptases and avoids strand-switching artifacts. dcDNA-seq, on the other hand, captures both strands, but is limited by the requirement of strand-switching for full-length cDNA synthesis. **(b)** Stacked barplot showing the biotype percentage of reads generated by each RT enzyme in either magnesium (MgCl_2_) or manganese (MnCl_2_) buffer. Biotypes include non-coding RNAs (ncRNA), ribosomal RNAs (rRNA), small nucleolar RNAs (snoRNA), mitochondrial RNAs (mtRNA), transfer RNAs (tRNA), spike-in RNAs (Sequins), and structured RNA IVTs (IVT) **(c,d)** Barplots (left panels) showing mismatch frequency across Watson-Crick (WC)-neutral and WC-disrupting bases for each RT enzyme in MgCl_2_ (c) and MnCl_2_ (d) buffers. The bar colors represent different base types: adenine (A), thymine (T), cytosine (C), guanine (G), and reference bases (gray). Line plots (right panels) depict RT drop-off scores as a function of position in the 25s rRNA, with WC-disrupting modifications (m^1^A, m^3^U, m^3^C) highlighted. **(e-f)** Barplot showing read length (nt) distribution of reads mapping to Sequins (left panel) and rRNAs (right panel), obtained from FIRST-seq data of the yeast total RNA sequencing using various reverse-transcriptases in magnesium and manganese buffers.

To address this gap, here we developed FIRST-seq (**Figure 2a**), a method that allows sequencing the first-strand cDNA products –including truncated RT products– transcriptome-wide, without the need for second strand synthesis or PCR amplification. Notably, FIRST-seq can be coupled to any reverse transcriptase of interest, as it does not require RT enzymes to leave a specific overhang at the 3’ end of the cDNA to perform second-strand synthesis. Moreover, it allows simultaneous capture of both RT drop-off information (**Figure S2a,b**) –which is typically lost in dcDNA-based methods– , in addition to basecalling errors (misincorporations, deletions and insertions) (**Figure S2c,d**), thus maximizing the captured signal. By contrast, we show that dcDNA-seq does not capture reverse transcription termination events induced by WC-disrupting modifications, limiting its suitability for RNA modification detection. Notably, we observed that dcDNA-seq did show sharp variations in coverage at specific positions along the yeast 25s rRNA, but none of these positions overlapped with annotated modified sites (**Figure S2a**).

### Systematic benchmarking of RT drop-off and mismatch signatures across RT enzymes and buffer conditions

FIRST-seq allows the use of distinct reverse transcriptases by bypassing the strand-switching step (**Figure 2a**). Here, we exploited this feature to examine which RT enzymes and buffer conditions might enhance the detection of chemical-induced and natural RNA modifications in the form of systematic basecalling errors and/or RT drop-offs. To this end, we systematically evaluated the performance of 8 different RT enzymes (Protoscript, Superscript II (SS2), Superscript III (SS3), Superscript IV (SS4), Maxima RT H Minus (Maxima), Thermostable Group II Intron RT (TGIRT), Induro RT (Induro) and Marathon RT (Marathon)) (see also **Table S3**) on a mix of RNAs (*S. cerevisiae* total RNA and several *in vitro*-transcribed RNAs: *B. subtilis* guanine riboswitch, *B. subtilis* lysine riboswitch, *Tetrahymena* ribozyme and sequin RNAs) **(Figure S3a,b).** The RNA mix was reverse transcribed using different RT enzymes and buffer conditions (MgCl2 or MnCl2, see also *Methods*), and the cDNA products were sequenced using FIRST-seq, followed by basecalling and mapping of the reads (see *Methods*).

First, we assessed how the choice of enzyme and buffer affected the composition of the sequencing library. We found that the proportion of recovered RNA biotypes varied significantly across conditions. Notably, the use of a MnCl2 buffer resulted in an increased proportion of recovered tRNAs for most enzymes, suggesting that manganese helps RTs overcome the extensive modifications and structures in tRNAs that typically cause stalling (**Figure 2b**, **Table S4**).

A highly desirable feature of RT enzymes for modification mapping is the introduction of misincorporations upon encountering a modified base. We systematically examined the ability of each RT enzyme to introduce basecalling ‘errors’ at known modified sites and found that the use of a MnCl2 buffer led to a striking and universal increase in mismatch rates for all eight enzymes tested (**Figure 2c-d**, left panels; **see also Table S5**). These results unequivocally demonstrate that manganese-based buffers are the superior choice for enhancing the detection of RNA modifications using ‘mutational profiling’ (MaP) methods. Interestingly, the increased error rate was not confined to Watson-Crick-disrupting modifications; we also observed a higher mismatch frequency at WC-neutral sites (**Figure 2c-d**), a feature that could be exploited to detect a broader range of RNA modifications.

In addition to error rates, we investigated the effect of buffer conditions on RT processivity by analyzing RT drop-off patterns on highly modified rRNAs. As expected, MnCl2 buffers consistently decreased premature RT termination at known WC-disrupting modification sites for all enzymes tested (**Figure 2c-d**, right panels, see also **Figure S3c-d**). However, this reduction in specific drop-offs did not uniformly translate to an increase in median read length across the library. An analysis of both synthetic sequin RNAs (**Figure 2e**) and endogenous rRNAs (**Figure 2f**) revealed that median read lengths were often shorter or did not significantly increase in MnCl2 conditions. This suggests that while manganese helps bypass specific blocking modifications, it may also contribute to general RNA fragmentation, a known effect of manganese ions. Nevertheless, the impact of manganese on generating full-length products was highly enzyme-dependent, particularly for shorter, structured non-coding RNAs. For instance, SS2 showed a marked increase in the proportion of full-length tRNAs and ncRNAs when used with a MnCl2 buffer, whereas this benefit was not observed for other enzymes like SS3 or TGIRT (**Figure S3e-f)**. This highlights that the overall efficiency of generating full-length cDNAs from challenging templates is determined by the specific interplay between the divalent cation and the reverse transcriptase. Notably, MnCl2 buffer also increased the basecalling error at WC-neutral modified sites (**Figure 2c-d**, see also **Figure S4a**), a feature that could be potentially further exploited to detect RNA modifications that otherwise cannot be detected via RT errors [5,9].

Collectively, these results highlight a key trade-off: manganese buffers significantly enhance the error signatures and read-through needed for modification detection, but this comes at the cost of potentially reduced overall read length due to RNA fragmentation. The optimal choice is therefore determined by the specific interplay between the divalent cation and the reverse transcriptase for the RNA templates of interest.

### Integrative analysis of RT drop-off and mismatch signatures identifies distinct enzymatic profiles and guides enzyme selection

To systematically compare the performance of reverse transcriptase (RT) enzymes, we performed principal component analysis (PCA) on four key features captured by FIRST-seq: i) biotype proportions, ii) mismatch frequency at modified sites, iii) mismatch base frequency at modified sites, and iv) RT drop-off rates at modified sites (**Figure 3a-e)**. This integrated analysis revealed that enzymes cluster into distinct families. Clustering by biotype revealed that viral RT enzymes (PS2 and SS2; SS4 and Maxima) and Group II intron RTs (Induro, TGIRT, and Marathon) generally clustered together, reflecting their ability to reverse transcribe molecules with similar features (**Figure 3a**). The exception to this trend was SS3, which clustered independently from the rest of viral RT enzymes under MgCl₂ conditions, and clustered with Group II intron RTs under MnCl₂ conditions.

**Figure 3.**
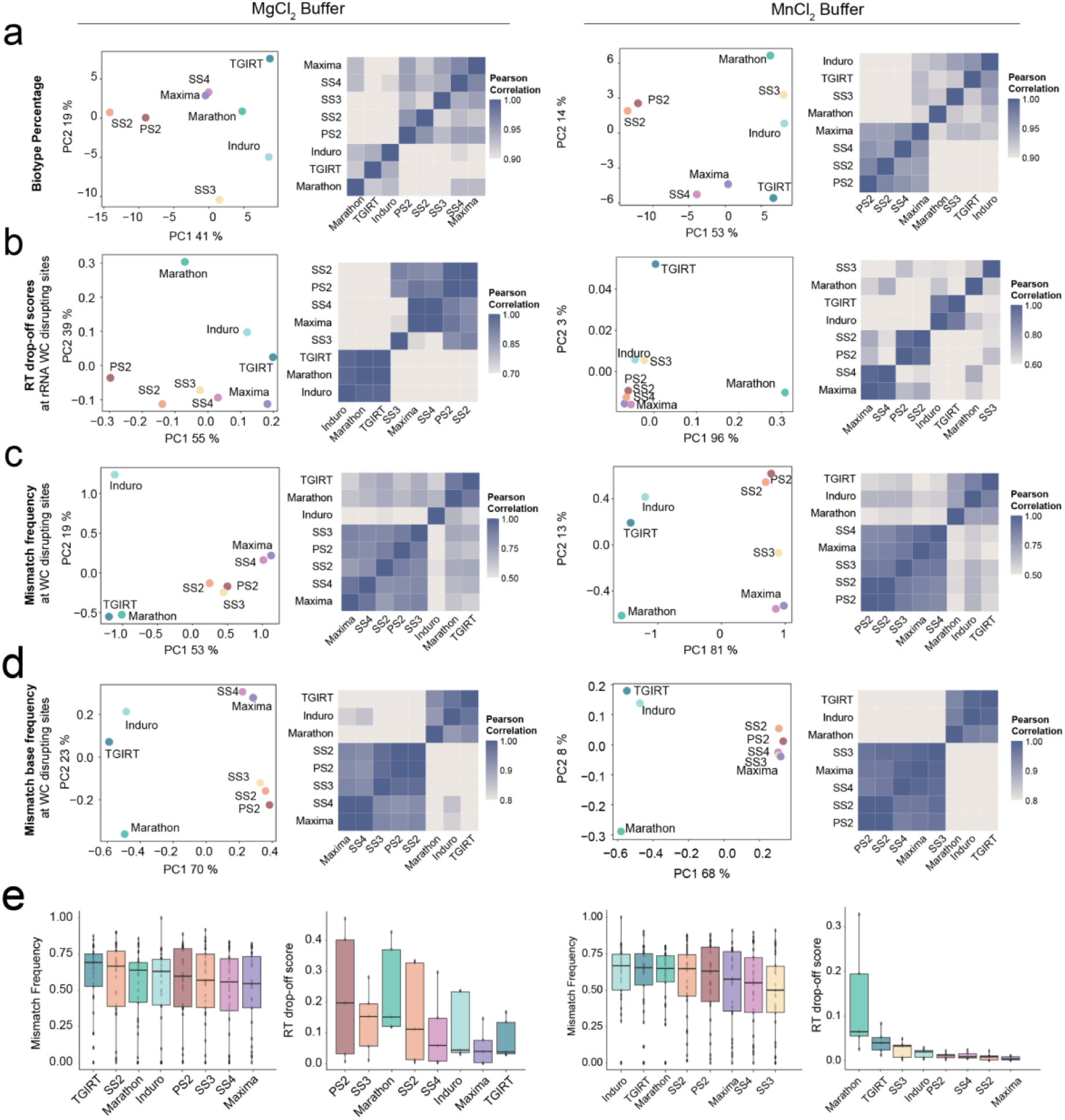
Signatures and performance metrics of reverse transcriptase enzymes in MgCl_2_ and MnCl_2_ buffers on the RNA mix sample used for benchmarking. **(a)** PCA plots showing the first two principal components (PC1 and PC2) based on biotype percentages from reads generated by different RT enzymes in MgCl_2_ (left) and MnCl_2_ (right) buffers. Each point represents an RT enzyme. Heatmap showing Pearson correlation coefficients of biotype percentages across RT enzymes in MgCl_2_ (left) and MnCl_2_ (right) buffers. **(b)** PCA plot showing the first two principal components (PC1 and PC2) of RT drop-off scores at rRNA Watson-Crick-disrupting (WC-disrupting) sites in MgCl_2_ (left) and MnCl_2_ (right) buffers. Heatmap showing Pearson correlation coefficients of RT drop-off scores at rRNA WC-disrupting sites of RT enzymes in MgCl_2_ (left) and MnCl_2_ (right) buffers. **(c)** PCA plots showing the first two principal components (PC1 and PC2) based on mismatch frequencies at WC disrupting sites for each RT enzyme in MgCl_2_ (left) and MnCl_2_ (right) buffers. Heatmap showing Pearson correlation coefficients of mismatch frequencies at WC-disrupting sites of RT enzymes in MgCl_2_ (left) and MnCl_2_ (right) buffers. **(d)** PCA plots showing the first two principal components (PC1 and PC2) based on mismatch base frequencies at Watson-Crick (WC) disrupting sites for each RT enzyme in MgCl_2_ (left) and MnCl_2_ (right) buffers. Heatmap showing Pearson correlation coefficients of mismatch base frequencies at WC-disrupting sites of RT enzymes in MgCl_2_ (left) and MnCl_2_ (right) buffers. **(e)** Boxplots of mismatch frequency and RT drop-off score for each RT enzyme in MgCl_2_ (left) and MnCl_2_ (right) buffer. For the mismatch frequency boxplots, each dot represents an m1A or m3C at rRNAs or tRNAs. For the RT drop-off score boxplots, each dot represents a WC-disrupting modification at rRNAs.

A key driver of this separation was the distinct error profiles of the enzyme families. PCA based on mismatch frequencies at WC-disrupting sites (**Figure 3c**) and mismatch base frequencies (**Figure 3d**) showed separate clustering of group II intron RTs and viral RT enzymes, suggesting that these two RT enzyme families produce dissimilar error patterns upon encountering a modified site. At known m¹A and m³C sites, the Group II intron RT enzymes (TGIRT, Induro, and Marathon) consistently produce the highest mismatch frequencies. This distinction is apparent in magnesium buffer but becomes even more pronounced in manganese buffer, where all enzymes exhibit an increased error rate (**Figure 3e**). To directly compare these signatures, we plotted the mismatch frequency of each RT against that of SS3 (**Figure S4b**). This analysis revealed that while manganese generally increases mismatch rates, the magnitude of this effect varies significantly between enzymes. In MnCl₂, mismatch frequencies for

Group II intron RTs (TGIRT, Induro and Marathon) increased far more dramatically than for SS3. This highlights a differential sensitivity to manganese, where Group II intron enzymes become exceptionally error-prone at modified sites compared to a viral RT like SS3. This behaviour is a key feature that can be exploited to distinguish specific modification types.

Enzyme processivity also contributed to the separation. PCA based on RT drop-off scores at WC-disrupting sites showed that Group II intron RTs clustered separately from M-MLV RTs (**Figure 3b**). A quantitative analysis of RT drop-off scores at Watson-Crick-disrupting sites within rRNA shows that manganese dramatically improves enzyme processivity. In magnesium, most enzymes exhibit high drop-off scores, indicating frequent premature termination. However, in a manganese buffer, these RT drop-off scores are universally minimized, signifying that the enzymes can more effectively read through blocking modifications (**Figure 3e**). Marathon RT is a notable exception, retaining a higher tendency for drop-off compared to other enzymes and clustering separately from other Group II intron RTs, likely due to its requirement for higher manganese concentration in order to minimize the drop-off events (**Figure 3b, e**) [9].

Based on these observations, we selected Induro and SS2 reverse transcriptases in a MnCl₂ buffer for DMS probing coupled with FIRST-seq (DMS-FIRST-seq). This selection was guided by the following criteria: i) a Mn^2+^-based buffer was chosen over Mg^2+^-based buffer, as it was found to maximize mismatch signatures while minimizing RT drop-off (**Figure 2c-d** and **3e**); ii) RT enzymes that produced stronger mismatch signatures were prioritized (i.e., Induro, TGIRT, Marathon RT and SS2 RT) (**Figure 3e**); iii) RT enzymes exhibiting high RT drop-off under Mn^2+^ buffer conditions were excluded (i.e., Marathon RT) (**Figure 3e**, right panel).

### DMS-FIRST-seq accurately predicts RNA structure *in vitro* and *in vivo*

We next assessed whether DMS-FIRST-seq would accurately recapitulate known RNA secondary structures. To this end, we used DMS-probed synthetic RNAs (*B. subtilis* lysine riboswitch and *Tetrahymena* ribozyme) as well as in *in vivo* DMS-treated cells (*E. coli* and human MDA-MB231, see ***Methods***). DMS-treated samples were prepared both using DMS-FIRST-seq and commercial dRNA-seq library preparation methods, to obtain a direct comparison of the two methods on the same treated samples.

We first confirmed that DMS treatment caused increased mismatch errors, which were captured both by DMS-FIRST-seq and dRNA-seq (**Figure 4a–b**; see also **Table S7**). Errors were not randomly distributed in the case of DMS-FIRST-seq, but rather were present largely at adenosine (A) and cytosine (C) residues. By contrast, dRNA-seq showed increased errors at all four bases **(Figure 4c-d)**. Consistent with prior work, cytosine base positions exhibited higher error rates than other bases in both FIRST-seq and dRNA-seq [43].

**Figure 4.**
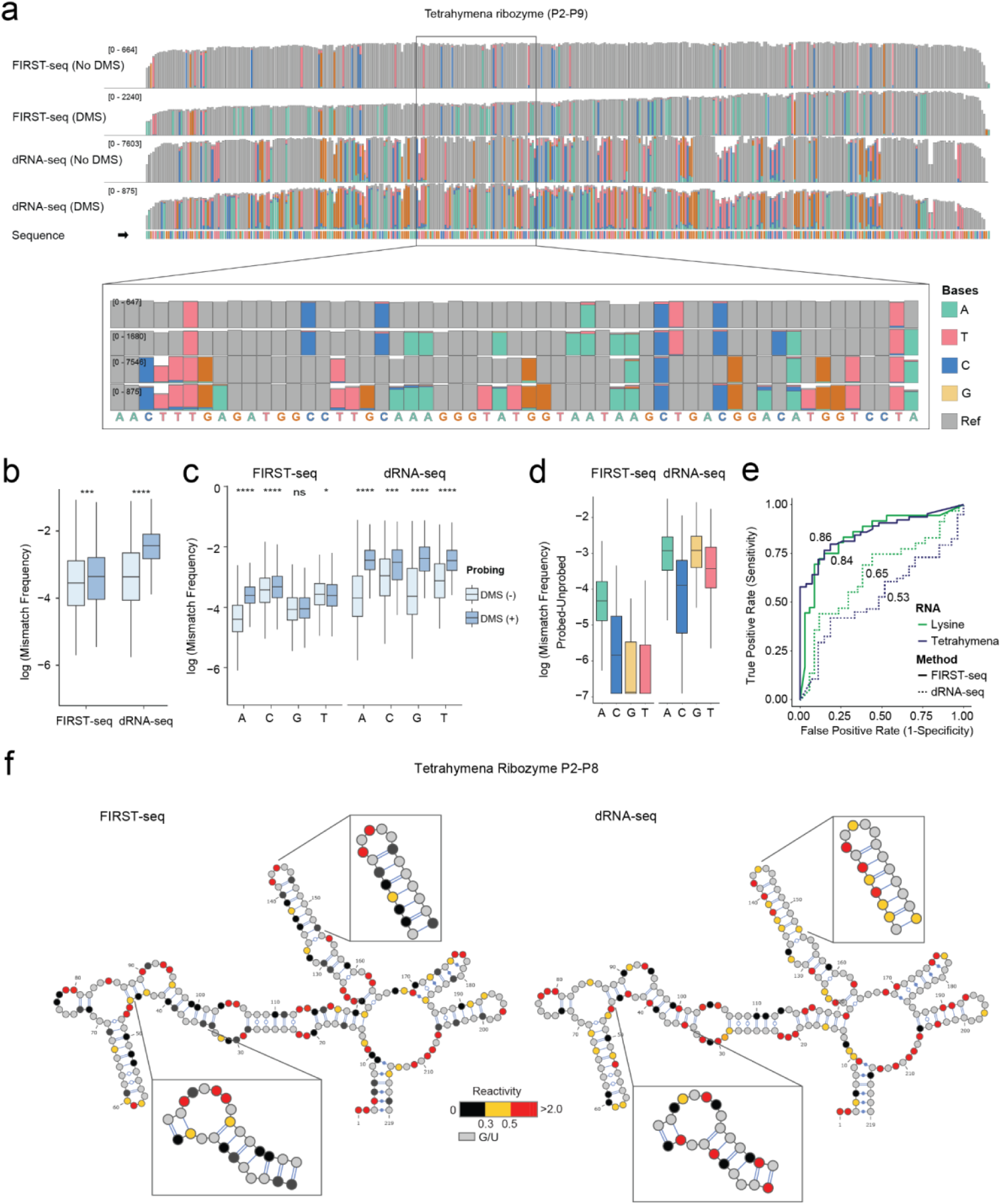
Structure assessment using *in vitro* DMS probing coupled to FIRST-seq. **(a)** Integrated Genomics Viewer (IGV) coverage track snapshots show FIRST-seq and dRNA-seq data of the unprobed and DMS probed Tetrahymena riboswitch. The zoomed box depicts a specific region within the RNA. Colored bars on top of the barplots indicate the modified positions. Mismatch frequency threshold for the coverage tracks is 0.02. Boxplots showing overall mismatch frequency change between unprobed and DMS probed RNAs sequenced using FIRST-seq and dRNA-seq. **(b)** Boxplots showing overall mismatch frequency change between unprobed and DMS probed RNAs sequenced using FIRST-seq and dRNA-seq.**(c)** Boxplots showing base-specific mismatch frequency change between unprobed and DMS probed RNAs sequenced using FIRST-seq and dRNA-seq. **(d)** Paired base-specific mismatch frequency difference between unprobed and DMS probed RNAs sequenced using FIRST-seq and dRNA-seq. **(e)** ROC curves comparing the True positive rate (Sensitivity) and False positive rate (1-Specificity) of FIRST-seq and dRNA-seq methods at agreeing with the known secondary structures of Lysine riboswitch and Tetrahymena ribozymes. Values shown on the lines are Area Under the Curve (AUC) values, assessing how well each method’s DMS reactivity correlates with the structures. **(f)** Secondary structure models with overlaid base reactivities for Tetrahymena ribozyme P2-P8 domain, obtained using FIRST-seq (left panel) and dRNA-seq (right panel). Structures have been generated using VARNA GUI version 1.0. Non-reactive bases (G/U) are colored in gray. For the boxplots, the box extends from the first quartile to the third quartile of the data, with a line at the median. The whiskers extend from the box to the farthest data point lying within 1.5x the interquartile range from the box. Statistical analyses were performed using the Kruskal–Wallis test (p > 0.05:ns, p ≤ 0.05:*, p ≤ 0.01:**, p ≤ 0.001:***, p ≤ 0.0001:**** ).

We then examined the accuracy of the both methods at recapitulating known RNA structures. To this end, DMS reactivity scores were calculated by subtracting the mismatch error rates between *in vitro* DMS-probed and unprobed samples on A and C positions (see also ***Methods*** and **Table S8**). Our results showed that DMS-FIRST-seq reactivities were highly consistent with known secondary structures, achieving AUC values of 0.84 and 0.86, for *B. subtilis* riboswitch and *Tetrahymena* ribozyme, respectively **(Figure 4e**). By contrast, DMS reactivity scores obtained from dRNA-seq showed lower AUC values (0.65 and 0.53, respectively) for the equivalent set of RNA molecules **(Figure 4e**). Visual inspection of DMS reactivity scores on the secondary structure of the *Tetrahymena* ribozyme P2–P8 domain confirmed that highly reactive bases corresponded to unpaired positions (**Figure 4f**). Together, these results confirmed that DMS-FIRST-seq outperformed dRNA-seq for accurate, base-resolution RNA structure mapping.

Next, we evaluated the performance of DMS-FIRST-seq *in vivo*. DMS incorporation was validated using LC-MS/MS, by quantifying the levels of DMS-induced modifications (m^1^A, m^3^C and m^7^G) upon increasing DMS concentrations **(Figure 5a,b).** We first examined the mismatch distribution in transcripts that showed high sequencing coverage, such as *Homo sapiens* RN7SL1 **(Figure 5c)** and *Escherichia coli* 16s ribosomal RNA **(Figure S4a),** finding that errors were significantly increased in DMS-treated samples compared to untreated controls **(Figure S4b-c,** see also **Tables S9-10),** as expected. Base-specific mismatch analysis revealed that errors were enriched at A and C positions, consistent with the specificity of DMS-induced modifications **(Figure 5d-e**, see also **Figure S4d-e)** and our results *in vitro* (**Figure 4**). High AUC values were obtained across distinct transcripts with known RNA structures, both for *E. coli* and human samples, confirming the ability of FIRST-seq to capture DMS-induced errors accurately also *in vivo* (**Figure 5f**, see also **Table S11**). Positions with high DMS reactivity scores mainly corresponded to unpaired positions, thus demonstrating its ability to capture RNA structural information (**Figure 5g)**.

**Figure 5.**
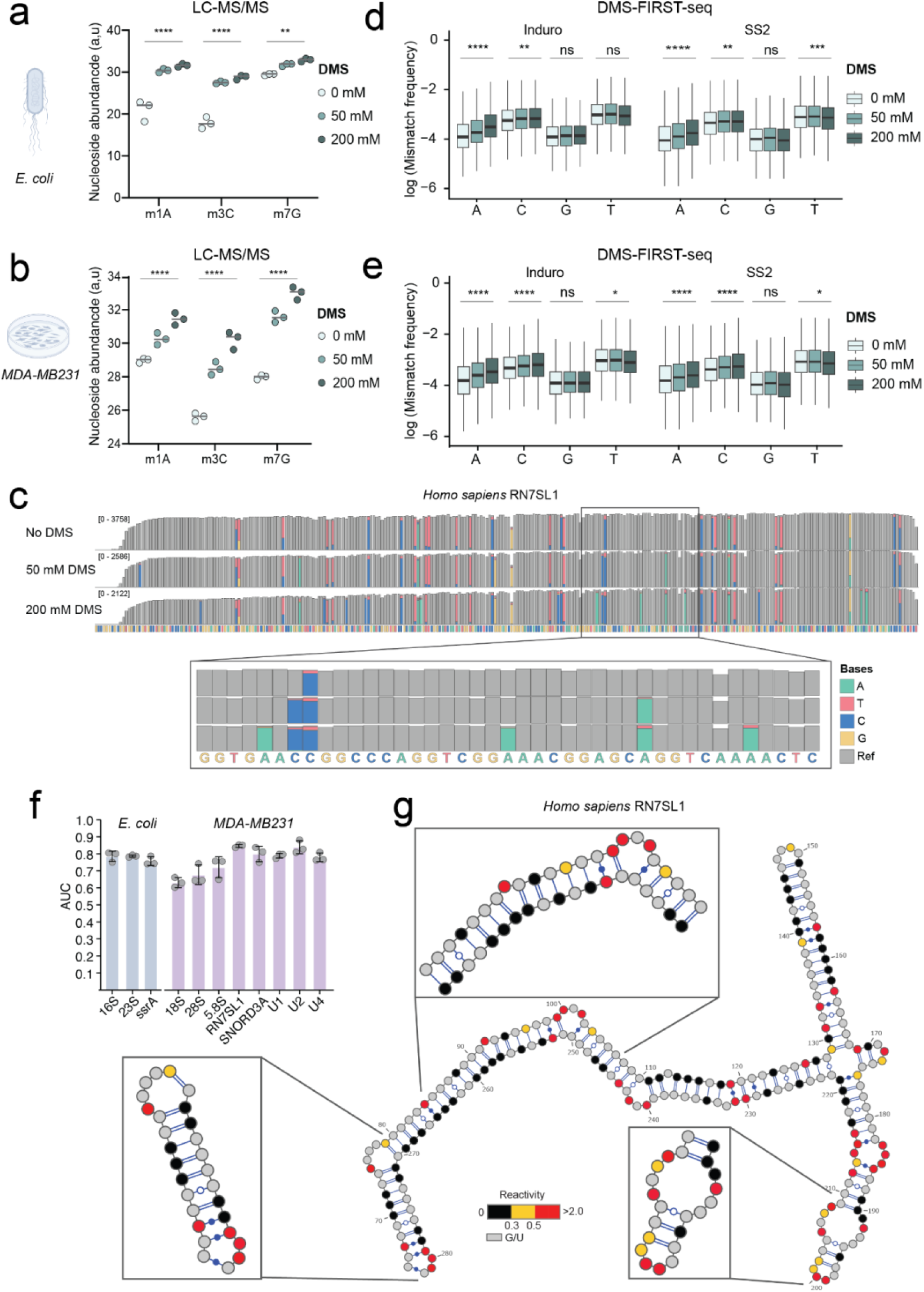
Structure assessment using *in vivo* DMS probing coupled to FIRST-seq in *E.coli* and *MDA-MB231* cell lines. **(a-b)** Dotplots showing nucleoside abundances (a.u.) of DMS induced base modifications m^1^A, m^3^C and m^7^G obtained using LC-MS/MS in the form of log2(Area Normalized to unmodified AUGC bases) for 0, 50 and 200 mM DMS probed *E.coli* **(a)** and MDA-MB231 **(b)** total RNAs. Each dot represents one replicate. **(c)** Integrated Genomics Viewer (IGV) coverage track snapshots show FIRST-seq data of the unprobed and DMS probed *H. sapiens* RN7SL1 RNA. The zoomed box depicts a specific region within the RNA. Mismatch frequency threshold for the coverage tracks is 0.05. **(d-e)** Boxplots showing base-specific mismatch frequency of the E.coli **(d)** and MDA-MB231 **(e)** total RNA probed with different concentrations of DMS prior to sequencing with FIRST-seq using Induro and SS2 enzymes. **(f)** AUC values calculated for showing the agreement between data obtained using FIRST-seq and known structure of various RNAs in *E.coli* and *H.sapiens* (MDA-MB231) cells treated with DMS. Scores were computed using A/C positions only, with reactivities defined as mismatch frequencies in 200 mM after subtracting the background in untreated cells. **(g)** Secondary structure model with overlaid base reactivities for *Homo sapiens* RN7SL1 RNA, obtained by using error differences between 200 mM DMS and 0 mM DMS treated RNAs. Structures have been generated using VARNA GUI version 1.0. Non-reactive bases (G/U) are colored in gray. For the boxplots, the box extends from the first quartile to the third quartile of the data, with a line at the median. The whiskers extend from the box to the farthest data point lying within 1.5x the interquartile range from the box. Statistical analyses were performed using the Kruskal–Wallis test (p > 0.05:ns, p ≤ 0.05:*, p ≤ 0.01:**, p ≤ 0.001:***, p ≤ 0.0001:**** ).

Finally, we compared the performance of DMS-FIRST-seq depending on the RT enzyme (Induro or SS2) by determining the agreement between DMS reactivities and known RNA secondary structures, for each enzyme. Our results revealed that Induro RT significantly outperformed SS2 (with Mn buffer) for *in vivo* RNA structure mapping at higher DMS concentrations (200mM) (**Figure S4f**).

Altogether, our results demonstrate that DMS-FIRST-seq enables accurate and transcript-specific RNA structure mapping *in vivo,* and that Induro RT coupled with the use of Mn-based buffers constitutes an optimal solution to capture RNA structure information when using long-read sequencing technologies.

### Detection of endogenous m^1^A modifications using FIRST-seq

The presence and distribution of m^1^A modifications in human cells has been a topic of active debate [44,45]. Reported estimates of m^1^A abundance vary widely, ranging from a few dozen [3], to several hundred [46] and even thousands [47] of putative m^1^A sites across the transcriptome. A major contributor to these discrepancies has been the use of m^1^A antibodies, which were subsequently found to cross-react with m^7^G caps [48]. Consequently, a fully reconciled human m^1^A map is still unavailable, largely because simple and robust methods for orthogonal validation of predicted m^1^A sites are lacking.

Here, we investigated whether FIRST-seq could serve as an antibody-independent method for orthogonal validation of m¹A-modified sites. We first systematically analyzed mismatch signatures at annotated m¹A sites in rRNAs and tRNAs. This revealed that Group II intron RTs, including TGIRT, Induro, and Marathon, consistently generated high mismatch rates at established m¹A sites **(Figure 6a,b)**. In contrast, viral RTs displayed lower mismatch frequencies, as well as distinct base misincorporation patterns (**Figure 6a,b**, see also **Figure S5**). Based on these observations, we hypothesized that employing two RT enzymes with divergent error patterns would improve the identification of m^1^A sites compared to the use of a single enzyme –typically TGIRT in previous studies [3,17,46]–. Accordingly, we selected TGIRT (to maintain comparability to previous work) and SS3 (the RT displaying the most distinct error signatures relative to TGIRT) as our enzyme pair, as they produce highly divergent mismatch signature profiles (**Figure 6a**, see also **Figure S6**).

**Figure 6.**
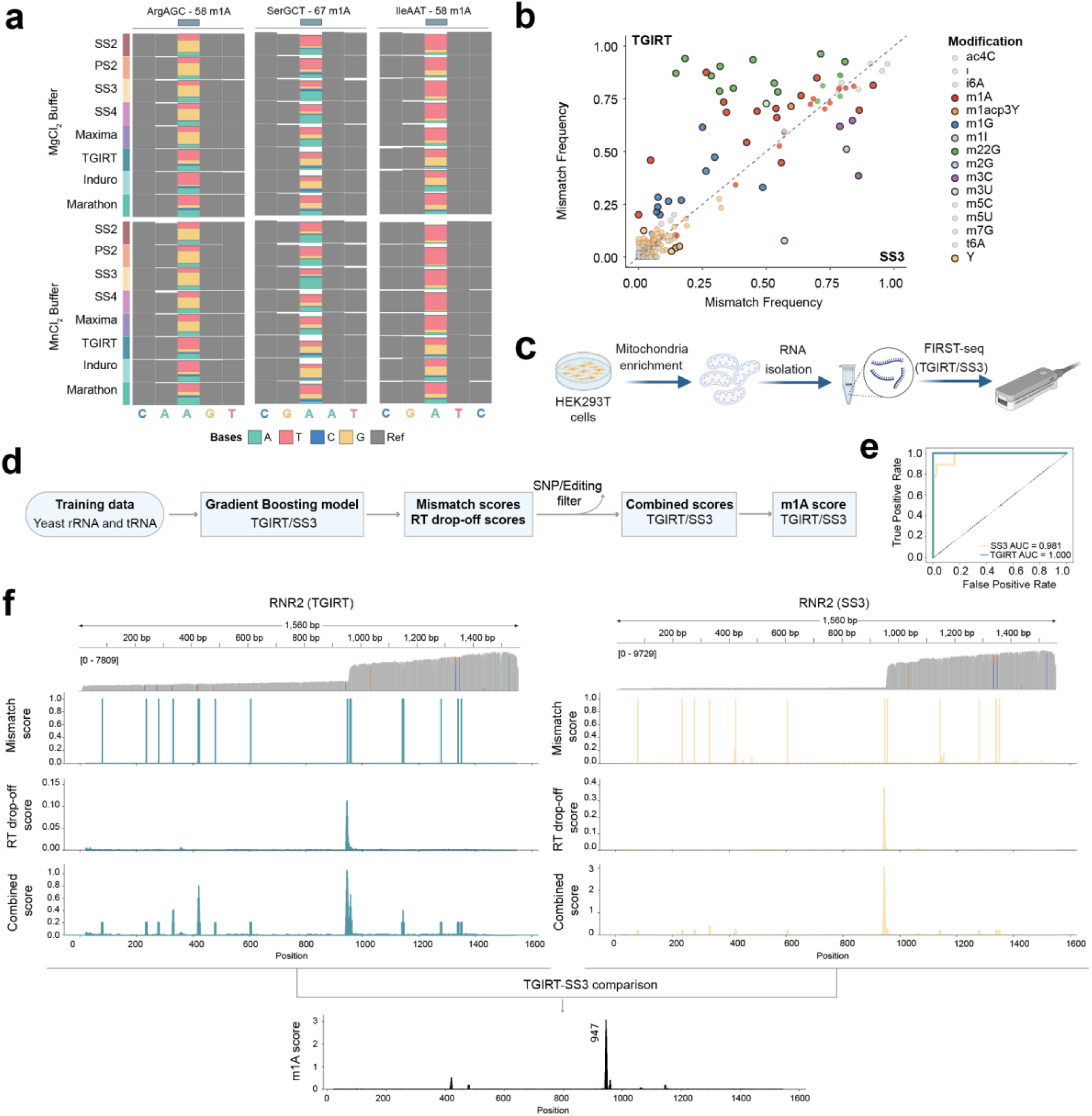
Detection of endogenous m¹A modifications using FIRST-seq and a comparative enzymatic approach. **(a)** Mismatch frequency profiles at known m¹A sites in three different yeast tRNAs (Arg-AGC, Ser-GCT, and Ile-AAT). The performance of eight different reverse transcriptases is compared under both magnesium and manganese buffer conditions. The color of the bars represents the base called at that position (A, T, C, G) or the reference base (Ref). Group II intron RTs (TGIRT, Induro, Marathon) in manganese buffer consistently yield the highest mismatch signals at m¹A sites. **(b)** Scatter plot comparing the mismatch frequency of various known RNA modifications in yeast when using TGIRT (y-axis) versus SS3 (x-axis) reverse transcriptases. Each dot represents a specific modification site. Note that N¹-methyladenosine (m¹A, red dots) shows a significantly higher mismatch frequency with TGIRT compared to SS3, indicating a distinct, enzyme-dependent signature. See also **Figure S5** for comparisons of mismatch frequency for other RT enzyme pairs. **(c)** Schematic of the experimental workflow for m¹A detection in human cells. Mitochondria were isolated from HEK293T cells, followed by total RNA extraction and the preparation of FIRST-seq libraries using two different enzymes, TGIRT and SS3. **(d)** Overview of the machine learning (ML) and data analysis pipeline for m¹A prediction. Separate Gradient Boosting (GB) models were trained on yeast tRNA and rRNA datasets for TGIRT and SS3. The trained models generate probability scores, which are combined with RT drop-off scores and filtered for SNPs/editing sites. A final differential m¹A score is computed to capture the enzyme-dependent signature. Mismatch scores were computed using the mismatch frequency and the base frequency mismatch type (see *Methods*). **(e)** Receiver operating characteristic (ROC) curves showing the performance of the trained Gradient Boosting classifiers for TGIRT and SS3 on held-out test data. The Area Under the Curve (AUC) values confirm high classification accuracy for both models. **(f)** Application of the m¹A detection pipeline to the human mitochondrial 16S rRNA (RNR2), which contains a known m¹A modification at position 947. The top panels show Integrated Genomics Viewer (IGV) snapshots for TGIRT and SS3 libraries. The subsequent plots show the individual m¹A probability scores, RT drop-off scores, and combined scores for each enzyme. The final differential score (TGIRT-SS3) reveals a single, prominent peak precisely at the known m¹A site at position 947. See also **Figure S7** for m^1^A scores along the ND5 mRNA.

We next applied FIRST-seq to identify putative m¹A sites in human HEK293T mitochondrial RNAs, using both TGIRT and SS3 reverse transcriptases (**Figure 6c**). To this end, we developed a computational pipeline that integrates both differential mismatch signatures (SS3 versus TGIRT) with RT drop-off information (**Figure 6d,e**). Briefly, a Gradient Boosting (GB) model was trained on known m¹A sites derived from yeast rRNA and tRNA FIRST-seq datasets, using per-site mismatch frequency and the identity of misincorporated bases as features, for reads generated either with SS3 or TGIRT. RT drop-off rates were subsequently calculated for each position and combined with mismatch features to derive a final per-site m¹A score. This score was calculated by comparing the differential signatures between the TGIRT and SS3 datasets (see also *Methods*).

Using this pipeline, we validated two mitochondrial m¹A sites in HEK293T cells: position 947 in the 16S rRNA (RNR2) (**Figure 6f**) and at position 1374 in the ND5 mRNA **(Figure S7)**. These correspond to two of the three mitochondrial m¹A sites reported in common by previous studies [3,49] **(Table S12**). Collectively, these results demonstrate that FIRST-seq enables antibody-independent orthogonal validation of m¹A sites.

### Guiding reverse transcriptase choice in FIRST-seq experiments

To provide a comprehensive and comparative view of how RT choice and buffer composition influence RNA modification detection in FIRST-seq, we compiled the performance of each RT enzyme and buffer conditions, separately for each RNA modification type **(Figure 7)**. Performance was assessed using four orthogonal readouts: (i) absolute mismatch frequency at modified positions (**Figure 7a**); (ii) RT drop-off at modified sites (**Figure 7b**); (iii) normalized read length across RNA biotypes (**Figure 7c**); and (iv) relative mismatch base frequency (**Figure S6**), reflecting the misincorporation preferences (see **Figure 6a** and **S8** for representative examples).

**Figure 7.**
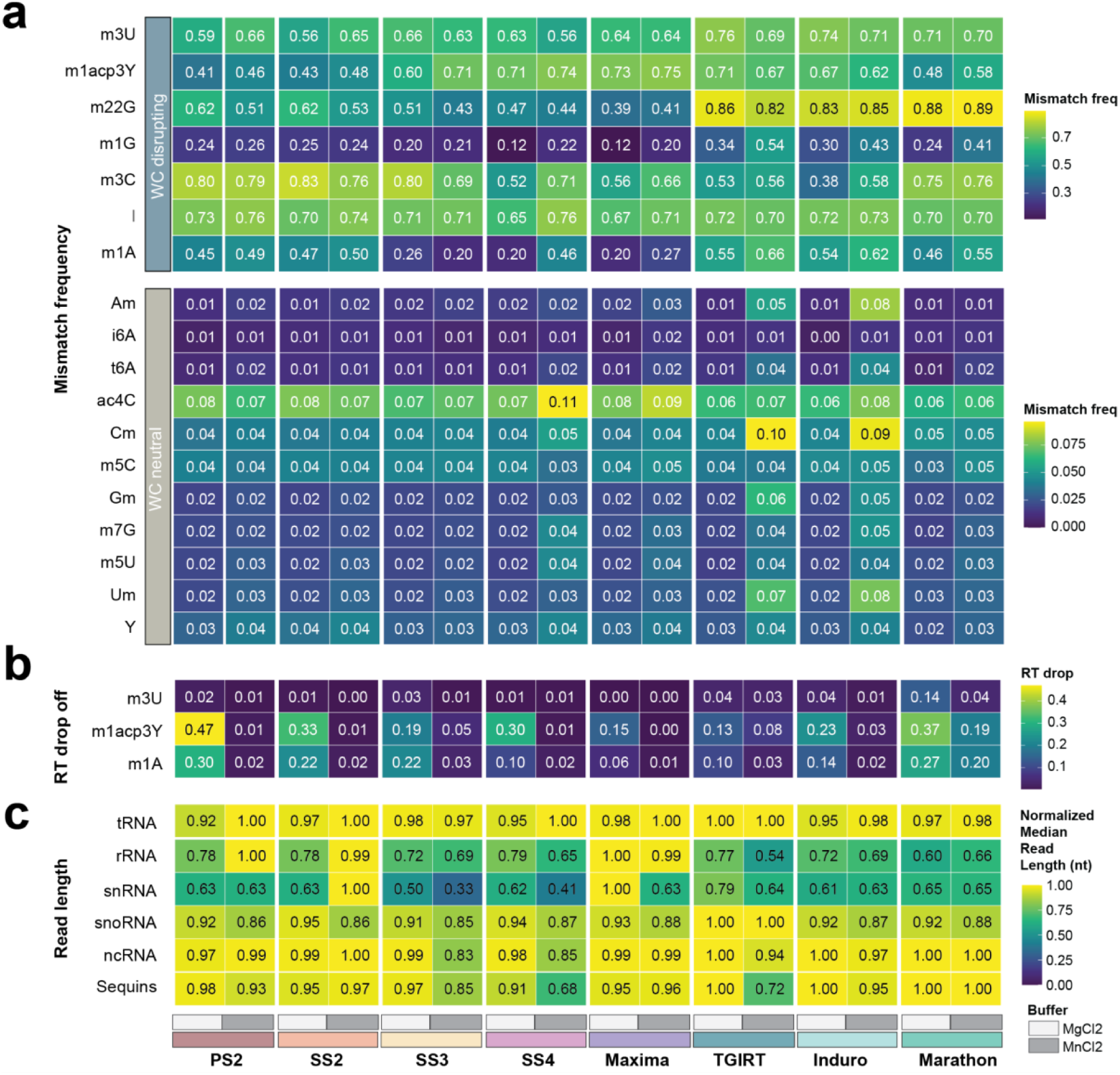
Integrated comparison of reverse transcriptase performance across FIRST-seq metrics. **(a)** Heatmap showing aggregated absolute mismatch frequencies from multiple modified positions in tRNAs and rRNA molecules, sequenced with each RT enzyme in either magnesium (MgCl_2_) or manganese (MnCl_2_) buffer. Color scale is separated for WC-disrupting and WC-neutral modifications for better visualization. **(b)** Heatmap showing aggregated absolute RT drop off scores from multiple modified positions in rRNA molecules, sequenced with each RT enzyme in either magnesium (MgCl_2_) or manganese (MnCl_2_) buffer. **(c)** Heatmap showing normalized median read length (nt) of reads mapped to different biotypes, sequenced with each RT enzyme in either magnesium (MgCl_2_) or manganese (MnCl_2_) buffer. The values have been scaled and normalized to allow for comparison across enzymes and biotypes simultaneously. A value of 1.00 is given to the enzyme and buffer condition that will show highest read length, for that given biotype.

Examination of these results revealed pronounced modification- and enzyme-specific behaviors. For example, m^2,2^G showed markedly elevated mismatch rates when reverse transcribed with Group II intron RTs (TGIRT, Induro, Marathon) (**Figure 7a**), consistent with increased misincorporation by these enzymes at WC-disrupting modifications [8,50]. Similarly, m¹A consistently produced stronger mismatch signatures with Group II intron RTs compared to viral RTs (**Figure 7a**). In contrast, m^3^C exhibited higher mismatch frequencies when reverse transcribed with viral RTs, underscoring that there is no ‘optimal’ RT enzyme for enhancing RT error signatures, but rather, that this response is modification-dependent.

Buffer composition further modulated modification detection, in agreement with previous works [51]. Notably, Nm modifications (Am, Cm, Gm, Um), which are typically not apparent as RT errors under Mg²⁺ buffer conditions, showed a clear increase in mismatch frequency in Mn²⁺-containing buffer (**Figure 7a**). A similar trend was observed for ac⁴C, which displayed substantially elevated mismatch frequencies under Mn2+ conditions; in this case, however, this phenomenon was most pronounced with the SSIV RT enzyme (**Figure 7a**).

Analysis of RT drop-off further highlighted enzyme-specific termination behaviors. Marathon RT exhibited pronounced RT drop-off across nearly all tested modifications, consistent with high sensitivity to base modifications but reduced RT read-through. While the use of Mn-based buffers overcame the RT drop-off for most RT enzymes tested, this effect was not observed for Marathon RT. (**Figure 7b**).

Systematic analysis of read length also revealed biotype-specific sensitivity to enzyme and buffer choice. Across most biotypes (tRNAs, snoRNAs, ncRNAs, and sequins), normalized median read length was consistently high across enzymes and buffers, indicating broadly comparable processivity of RT enzymes. By contrast, rRNAs and snRNAs showed the largest RT-dependent variation, with some enzyme–buffer combinations exhibiting significant reduction in read length relative to the best-performing condition within the same biotype (**Figure 7c**). Notably, Maxima RT displayed the highest overall processivity, yielding longest read lengths across biotypes, outperforming Group II intron RTs (TGIRT, Marathon RT and Induro RT).

Overall, these results demonstrate that no single reverse transcriptase performs optimally across all buffers, RNA biotypes and modification types. Rather, FIRST-seq enables application-driven enzyme selection, where different RT–buffer combinations are chosen depending on whether the goal is RNA structure probing or detection of a specific RNA modification.

## DISCUSSION

Nanopore direct RNA sequencing offers a simple and cost-effective platform to directly detect RNA modifications in native RNA molecules [30,38,39,52], including those introduced by chemical probing reagents [21]. Although modification-aware basecalling models have recently become available to detect certain RNA modifications at single-nucleotide and single-molecule resolution [31], models capable of identifying modifications induced by chemical probing reagents (e.g., m^1^A, m^3^C, and m^7^G upon DMS probing) are not yet available. In contrast, cDNA-based approaches enable the generation of single nucleotide resolution maps of unpaired probed positions (i.e., m^1^A and m^3^C following DMS treatment), and have the added advantage of being invisible to probed paired positions that do not contain RNA structure information (e.g., m^7^G modifications). Recently, methods such as Nano-DMS-MaP-seq have exploited this principle to obtain long-read RNA structures of viral transcripts using highly processive group II intron reverse transcriptases [22]. However, their dependence on gene-specific primers limits their applicability at the transcriptome-wide scale.

To overcome this limitation, we developed FIRST-seq, a platform that enables long-read, transcriptome-wide analysis through direct sequencing of the first cDNA strand. This approach eliminates the need for strand-switching, making it compatible with any reverse transcriptase (RT) and buffer system **(Figure 2a)**. This flexibility allowed systematic benchmarking of a panel of RT enzymes and buffer conditions, revealing that both enzyme selection and buffer composition critically impact performance, affecting RNA biotype recovery (**Figure 2b and 3a)**, mismatch frequencies (**Figure 2c**, and **3c-d**) and RT drop-off rates **(Figure 3b)**. Our benchmarking revealed a key trade-off: magnesium-based buffers promote high-fidelity synthesis, whereas manganese-containing buffers enhance both processivity and the error signatures required for modification profiling **(Figure 2e,f)**. Notably, although Group II intron RTs showed higher processivity at specific modified sites, certain viral RTs, such as SS2 and Maxima RT, yielded the highest proportion of full-length reads under optimized conditions, underscoring the importance of empirical optimization for transcriptome-wide applications.

We next assessed the applicability of the FIRST-seq platform in two distinct contexts. First, by coupling it with DMS probing (DMS-FIRST-seq), we generated high-accuracy RNA structure maps that showed strong concordance with known secondary structures , both *in vitro* (**Figure 4d-f**) and *in vivo* (**Figure 5f-g**). Second, we demonstrated its ability to detect and validate endogenous RNA modifications using a differential RT-signature-based pipeline, which we applied to identify m^1^A-modified sites in mitochondria (**Figure 6**). A key advantage of FIRST-seq is that it enables both single-enzyme and comparative, multi-enzyme RNA modification detection within the same experimental framework. While individual reverse transcriptases such as TGIRT provide high sensitivity for specific modifications (e.g., m¹A) [3], here we show that combining two RT enzymes with differential RT error signatures allows to achieve higher specificity than when using a single RT enzyme. This comparative strategy provides an intrinsic control against enzyme-specific artifacts, without requiring antibody enrichment [3] or chemical conversion steps [46].

It is important to note, however, that a key limitation of FIRST-seq lies in its reduced sensitivity for low-stoichiometry modified sites (e.g. <5%), as the method does not include pre-enrichment for modified RNA species. Consequently, it remains unclear whether the additional sites reported by previous studies –which were largely non-overlapping– represent false positives or modifications below the detection threshold of FIRST-seq.

Although we did not explicitly investigate the use of FIRST-seq for systematic identification and validation of other RNA modification types beyond m^1^A, our results suggest that this differential signature framework can be extended to additional modifications, such as m^2,2^G, m^1^G and m^3^C (**Figure 7** and **S5,6**), which also exhibit differential reverse transcription signatures when analyzed using specific RT enzyme pairs. To facilitate the use of RT combinations to detect distinct types of modifications, we provide a systematic evaluation of RT enzymes and buffers, separately for each RNA modification type **(Figure 7**, see also **Figure S6).** Through this analysis, we observe that different RNA modifications exhibit distinct —and in some cases opposing— sensitivities to RT enzyme, buffer composition, and termination behavior. For example, Group II intron RTs generate strong mismatch signatures at bulky base modifications such as m¹A and m²²G, whereas viral RTs tend to provide more uniform read-through but reduced sensitivity to certain modifications. Similarly, manganese buffers enhance mismatch detection at subtle modifications such as Nm residues, while also influencing read-length distributions in a biotype- and enzyme-dependent manner. Collectively, these results demonstrate that no single enzyme–buffer combination is optimal across all applications, and instead highlight FIRST-seq as a flexible platform that is compatible with any RT enzyme of interest, under flexible buffer conditions, enabling RNA structure and modification profiling.

Our results show that FIRST-seq can capture highly-structured and highly-modified RNA templates, such as tRNAs. We observed that the use of manganese buffer significantly increased the recovery of tRNAs, which are notoriously difficult to reverse transcribe due to their extensive post-transcriptional modifications and stable tertiary structures that are known to cause RT drop-off [53,54] **(Figure 2b** and **S3e)**. Thus, FIRST-seq can be used as a platform for epitranscriptomic studies, including those found in highly-structured, heavily modified RNA species.

In summary, FIRST-seq provides a flexible and powerful platform for transcriptome-wide RNA structure and modification analysis using long-read cDNA nanopore sequencing. By decoupling cDNA synthesis from the constraints of commercial library preparation kits, FIRST-seq enables the use of any reverse transcriptase and facilitates optimization of reaction conditions for diverse biological applications and RNA modification types.

## METHODS

### Generation of *in vitro* transcribed RNA molecules

The list of *in vitro* transcribed constructs used in this work can be found in **Table S13**, and the corresponding Fasta files can be found in https://github.com/novoalab/firstseq. *Tetrahymena* ribozyme sequence was obtained from IDT (Ref ID: 129935271). *Tetrahymena* ribozyme sequences were ordered as synthetic DNA constructs to IDT, flanked by BamHI and XhoI restriction sites and inserted into pIDTBlue plasmid. *B. subtilis* XPT_Guanine and Lysc_Lysine riboswitch sequences were ordered as synthetic DNA constructs to General Biosystems, flanked by DraI restriction sites, and inserted into a pUC57 plasmids. Sequin[55] plasmids, flanked by EcoRI-HF restriction sites (R2-63-3, R1-81-2, R1-103-1, R2-117-1) were kind gifts from Tim Mercer’s lab (University of Queensland). To obtain *in vitro* transcribed RNAs, maxipreps of each of the plasmids were prepared using Qiagen Plasmid Plus Maxi Kit (Qiagen, #12963). Each construct was linearized using specific restriction enzymes. XPT_Guanine and Lysc_Lysine riboswitch sequences were linearized using DraI (NEB, R0129S), the T*etrahymena* ribozyme sequence was linearized using BamHI (NEB, R0136S) and XhoI (NEB, R0146S) and sequins sequences were linearized using EcoRI-HF (NEB, R3101S). 20 ug of each plasmid was digested overnight at 37℃ in 500 ul reaction with 10U enzyme: 1 ug plasmid ratio. Digestion product was extracted using Phenol:Chloroform:Isoamyl alcohol (25:24:1). Digestion was confirmed with Agarose Gel Electrophoresis. The digested products of the synthetic *Tetrahymena* ribozyme, XPT_Guanine and Lysc_Lysine riboswitch sequences were *in vitro* transcribed using Ampliscribe™ T7-Flash™ Transcription Kit (Lucigen, ASF3507) following manufacturer’s instructions. Sequin sequences were *in vitro* transcribed using SP6 RNA Polymerase (NEB, M0207S) following manufacturer’s instructions. IVT RNA was then treated with Turbo DNase (Thermo, AM2238) (2 ul enzyme for 50 ul reaction of 200 ng/ul RNA) at 37°C for 15 minutes, with a subsequent Zymo RNA Clean & Concentrator-5 (Zymo, R1013) cleanup, following the manufacturer’s instructions. The quality of the *in vitro* transcribed (IVT) products as well as the addition of polyA tail to the synthetic constructs was assessed using Agilent 4200 Tapestation **(Figure S3a-b**). Concentration was determined using Qubit Fluorometric Quantitation. The purity of the IVT products was measured with NanoDrop 2000 Spectrophotometer.

### Yeast culturing and total RNA extraction

*Saccharomyces cerevisiae* (strain BY4741) was grown at 30°C in standard YPD medium (1% yeast extract, 2% Bacto Peptone and 2% dextrose). Cells were then quickly transferred into 50 mL pre-chilled falcon tubes, and centrifuged for 5 minutes at 3,000 g in a 4°C pre-chilled centrifuge. Supernatant was discarded, and cells were flash frozen. Flash frozen pellets were resuspended in 700 µL TRIzol (Thermo, 15596026) with 350 µL acid washed and autoclaved glass beads (425-600 µm, Sigma G8772). The cells were disrupted using a vortex on top speed for 7 cycles of 15 seconds (the samples were chilled on ice for 30 seconds between cycles). Afterwards, the samples were incubated at room temperature for 5 minutes and 200 µL chloroform was added. After briefly vortexing the suspension, the samples were incubated for 5 minutes at room temperature. Then they were centrifuged at 14,000 g for 15 minutes at 4°C and the upper aqueous phase was transferred to a new tube. RNA was precipitated with 2X volume Molecular Grade Absolute ethanol and 0.1X volume Sodium Acetate (3M, pH 5.2). The samples were then incubated for 1 hour at -20°C and centrifuged at 14,000 g for 15 minutes at 4°C. The pellet was then washed with 70% ethanol and resuspended with nuclease-free water after air drying for 5 minutes on the benchtop. Purity of the total RNA was measured with the NanoDrop 2000 Spectrophotometer. Total RNA was then treated with Turbo DNase (Thermo, AM2238) (2 ul enzyme for 50 ul reaction of 200 ng/ul RNA) at 37°C for 15 minutes, with a subsequent RNAClean XP bead cleanup.

### *In vitro* DMS probing of IVT RNAs

1 ug IVT RNA was denatured at 95℃ for 2 mins and cooled on ice for 2 mins. Then RNA was mixed with RNA Folding Buffer (10 mM Tris pH 8.0, 100 mM NaCl, 6 mM MgCl2) up to a volume of 50 uL and incubated at 30℃ for 30 mins. For probing, 1.25 uL DMS was added to the DMS probed RNA, and 1.25 uL water was added to the unprobed RNA, and mixes were incubated at 30℃ for 5 mins. Reaction was stopped by adding 100 uL of Quenching Solution (3M B-Mercaptoethanol, 508 mM NaAC, 2ul pellet paint for each reaction), and incubated for 5 min at room temperature. RNA was precipitated by adding 300 uL absolute ethanol 100%, and resuspended in 15 uL nuclease-free water. Probed and unprobed RNAs were then polyadenylated in vitro, to prepare them for nanopore sequencing. Poly(A) tailing was performed using *E. coli* Poly(A) Polymerase (NEB, M0276S) as per manufacturer’s instructions. Poly(A) tailed RNA was recovered using Zymo RNA Clean & Concentrator-5, following the manufacturer’s instructions. Concentration was determined using Qubit Fluorometric Quantitation. Purity of the IVT product was measured with a NanoDrop 2000 Spectrophotometer.

### In vivo DMS probing of *Escherichia coli*

For *E. coli*, a single colony of DH5alpha bacteria was picked and inoculated in LB broth, and grown for 16h at 37°C with shaking. Bacteria were then diluted to OD600 = 0.05 and allowed to grow for an additional 2h, until OD600 ∼ 0.4-0.5 (exponential phase). Dimethyl sulfate (DMS, cat. D186309, Merck), from a 1:6 dilution in ethanol (1.76 M), was directly added to the bacteria at the desired concentration. Bacteria were then shaken at 37°C for 2 min, after which DMS was immediately quenched by addition of 1 volume ice-cold DTT 1M. Bacteria were quickly pelleted by centrifugation at 17,000 g (4°C) for 1 min, the supernatant discarded, and pellets snap-frozen in liquid nitrogen.

### In vivo DMS probing of MDA-MB-231

MDA-MB-231 were grown in high glucose DMEM medium, supplemented with 10% fetal bovine serum, at 37°C and 5% CO2. After the second passage, cells at ∼80% confluence were probed by direct addition of DMS, from a 1:6 dilution in ethanol (1.76 M), to the culture medium. Cells were then incubated at 37°C for 2 min, after which DMS was immediately quenched by addition of 1 volume ice-cold DTT 1M. Cells were then quickly pelleted by centrifugation at 5,000 g (4°C) for 1 min, the supernatant discarded, and pellets snap-frozen in liquid nitrogen.

### RNA Isolation from *Escherichia coli and* MDA-MB-231 cells

For *E. coli*, bacterial pellets were resuspended in 62.5 μl Buffer R [20 mM Tris-HCl pH 8.0; 80 mM NaCl; 10 mM EDTA pH 8.0], supplemented with 100 μg/mL final Lysozyme (cat. L6876, Merck) and 20 U SUPERase•In™ RNase Inhibitor (cat. A2696, ThermoFisher Scientific), by thorough vortexing. After incubating samples at room temperature for 1 min, 62.5 μl Buffer L [0.5% Tween-20; 0.4% Sodium deoxycholate; 2 M NaCl; 10 mM EDTA] were added. Samples were then inverted 10 times and incubated at r.t. for 2 min, and on ice for 2 min. 1 mL ice-cold TRIzol™ Reagent (cat. 15596018, ThermoFisher Scientific) was then added, and samples were vortexed for 15 sec. RNA was extracted as per manufacturer protocol. For MDA-MB-231, cell pellets were directly resuspended in 1 mL TRIzol™ Reagent, incubated at 70°C for 5 min, and then RNA was extracted as per manufacturer protocol. Integrity of both *E. coli* and human DMS-probed total RNA was assessed using Agilent Tape Station (**Figure S9**).

### Mitochondrial RNA Isolation form HEK293 cells

Mitochondrial fractions were isolated from HEK293 cells as previously described [56]. Briefly, 3 × 10⁷ cells were pelleted and resuspended in 1.5 mL of ice-cold MitoPrep buffer (0.225 M mannitol, 0.075 M sucrose, 20 mM HEPES, pH 7.4, filtered through 0.22 μm filter and stored at 4°C) and transferred to a pre-chilled 5 mL glass/Teflon homogenizer. Cells were disrupted on ice with 30 strokes. The homogenate was transferred to a 1.5 mL microcentrifuge tube and centrifuged at 800 x g for 5 mins. The supernatant was collected, and the pellet was resuspended in another 1.5 mL of MitoPrep buffer and re-homogenized with 20 strokes on ice. This second homogenate was also centrifuged at 800 × g for 5 min. Supernatants from both homogenization steps were pooled and centrifuged again at 800 × g for 5 min to remove residual nuclei and debris. The resulting supernatant was centrifuged at 11,000 × g for 5 min to pellet crude mitochondria. Mitochondria pellet was resuspended in 30 μL of MitoPrep buffer (pH 7.4). Protein concentration of mitochondrial fraction was measured using a NanoDrop spectrophotometer. Then, 1 μL of protein sample was diluted in 19 μL of 0.6% SDS (w/v), and the absorbance was measured. Final protein concentrations were calculated by multiplying the measured values by a dilution factor of 20. Typically, 3 × 10⁷ cells yielded ∼800 μg of crude mitochondria. 200 μg of purified mitochondria (in 20 μl of MitoPrep buffer) was transferred to a microcentrifuge tube and kept on ice until RNA isolation.

Next, 200 μg of purified mitochondria (measured as described above and resuspended in 20 μL of MitoPrep buffer) was transferred to a microcentrifuge tube and kept on ice. Mitochondria were lysed by adding 100 μL of freshly prepared lysis buffer (1% SDS, 10 mM EDTA, 10 mM Tris-HCl, pH 7.4) and incubated at 70 °C for 5 min in a heat block, followed by cooling to RT. Protein digestion was performed by adding 1 μL of 1 mg/mL Proteinase K (Roche, cat. 3115836001) and incubating at 37 °C for 5 min.

Subsequent RNA extraction steps were performed in a chemical fume hood. Lysates were mixed with 400 μL of TRIzol™ Reagent (cat. 15596018, ThermoFisher Scientific) and 200 μL of 1-bromo-3-chloropropane (BCP) (Merck, cat. B9673), vortexed vigorously for 1 min, and centrifuged at 14,800 × g for 5 min at RT. The upper aqueous phase was transferred to a new microcentrifuge tube, mixed with an equal volume of isopropanol, and centrifuged at 21,000 × g for 10 min at 4 °C to pellet RNA. The RNA pellet was washed with 600 μL of ice-cold 75% ethanol, gently inverted several times, and centrifuged again at 21,000 × g for 5 min. The supernatant was carefully removed, and residual ethanol was minimized to avoid pellet loss. The RNA pellet was briefly air-dried in a laminar flow cabinet and subsequently treated with Turbo DNase (Thermo, cat. AM2238) according to the manufacturer’s instructions.

### Evaluation of agreement between DMS signal and structure

Agreement between DMS signal and reference structures for *in vitro* and *in vivo* RNAs was calculated using the *rf-eval* module of the RNA Framework[57]. Reference structures were obtained from the RNA Central database.

### DMS reactivity score calculation

For DMS score calculation, background-subtracted raw mutation rates were normalized via the *rf-norm* module of the RNA Framework [57], using the box-plot normalization approach (parameter: *-nm 3*).

### Nanopore direct RNA sequencing library preparation

The RNA libraries for direct RNA sequencing (SQK-RNA002) were prepared with the manufacturer’s protocol with some adjustments. Unprobed and DMS probed RNAs were polyadenylated using *E. coli* Poly(A) Polymerase, following the commercial protocol, before starting the library prep. RNA library for direct RNA sequencing (SQK-RNA002) was prepared following the ONT Direct RNA Sequencing protocol version DRS_9080_v2_revI_14Aug2019. Briefly 500 ng total of poly(A) tailed RNAs were ligated to RT adaptors (RTA) using concentrated T4 DNA Ligase (NEB, M0202T) and was reverse transcribed using TGIRT, without the heat inactivation step. The products were purified using 1.8X Agencourt RNAClean XP beads (Thermo, NC0068576) and washed with 70% freshly prepared ethanol. Then, RMX adapter, composed of sequencing adapters with motor protein, was ligated onto the RNA:DNA hybrid, and the mix was purified using 1X Agencourt RNAClean XP beads, washing with buffer twice. The sample was then eluted in elution buffer and mixed with RNA running buffer before loading onto a primed R9.4.1 flowcell and run on a MinION sequencer with MinKNOW acquisition software version v.3.5.5.

### Nanopore direct cDNA library preparation

The cDNA libraries for the direct cDNA sequencing (SQK-DCS109) were prepared following the direct cDNA Sequencing ONT protocol (DCB_9091_v109_revC_04Feb2019) and barcoded using the Native Barcoding Expansion kit (EXP-NBD104). Briefly, 50 ng poly(A) tailed RNA was mixed with 2.5 uL VNP oligo, 1 uL dNTPs (Thermo, R0192), and up to 11 uL nuclease-free water. After gently mixing by flicking and spinning down briefly, the mix was incubated at 65°C for 5 minutes and then placed on ice immediately. In a separate tube, 4 uL 5X RT Buffer, 1 uL RNase Inhibitor Murine (NEB, M0314S), 2 uL Strand-Switching Primer (SSP), 1 uL nuclease-free water was prepared and then two solutions were mixed. After incubating the mix at 42° C for 2 minutes, 1 uL Maxima H Minus Reverse Transcriptase (Thermo, EP0752) was added. Then the mix was incubated at 42°C for 90 minutes and inactivated by heating at 85°C for 5 minutes before moving to ice. RNAse Cocktail (Thermo Scientific, AM2286) was added to the mix in order to digest the RNA, and the mix was incubated at 37°C for 10 minutes. The reaction was then cleaned up using 0.8X AMPure XP Beads (Agencourt, A63881). Second-strand synthesis was performed by mixing the 20 uL cDNA sample with 25 uL 2X LongAmp Taq Master Mix (NEB, M0287S), 2 uL PR2 Primer (PR2), and 3 uL nuclease-free water. The reaction was incubated at 94 °C for 1 min, 50 °C for 1 min, 65 °C 15 mins, and placed on ice. The reaction was then cleaned up using 0.8X AMPure XP Beads. End-repair and dA-tailing of the double-stranded DNA was achieved by mixing the 20 uL DNA sample with 30 uL nuclease-free water, 7 uL Ultra II End-prep reaction buffer and 3 uL Ultra II End-prep enzyme mix (NEB, E7546S), and incubating this mix at 20° C for 5 minutes and 65° C for 5 minutes. After incubating the reaction on ice for 1 min, it was cleaned up using 0.8X AMPure XP Beads. End-prepped DNA was then ligated to the Native Barcode (NB) for multiplexing with other libraries. The ligation was performed by mixing 22.5 uL end-prepped DNA, 2.5 uL Native Barcode (NB), 25 uL Blunt/TA Ligation Master Mix (NEB, M0367S) and incubating the mix at room temperature for 10 mins. The reaction was then cleaned up using 0.8X AMPure XP Beads. Once all the barcoded libraries were pooled after elution, the ligation of the sequencing Adapter Mix (AMX) was performed by mixing the 65 uL pooled barcoded DNAs (up to 200 fmol), 5 uL AMX, 20 uL 5X NEBNext Quick Ligation Reaction Buffer (NEB,B6058S) and 10 uL Quick T4 DNA Ligase (NEB,M2200S), and incubating the mix at room temperature for 10 mins. The reaction was cleaned up using 0.8X AMPure XP beads, using WSB Buffer for washing. The sample was then eluted in Elution Buffer (EB) and mixed with Sequencing Buffer (SQB) and Loading Beads (LB) prior to loading onto a primed R9.4.1 flowcell. The library was run on MinION flowcell with MinKNOW acquisition software version v.3.5.5.

We should note that the direct cDNA kit (SQK-DCS109) used in this work is now deprecated; however, equivalent direct cDNA libraries can be built using the SQK-LSK114 kit, which is compatible with the current R10 flowcells, using the following protocol (https://nanoporetech.com/document/ligation-sequencing-v14-direct-cdna-sequencing).

### FIRST-seq library preparation

Before starting the library preparation, 2 µL of 10 µM VNP (/5Phos/ACTTGCCTGTCGCTCTATCTTCTTTTTTTTTTTTTTTTTTTTVN) and 2 µL of 10 µM Comp_DNA (GAAGATAGAGCGACAGGCAAGTA) were mixed with 1 µL 0.1 M Tris pH 7.5, 1 µL 0.5 M NaCl, and 4 ul RNase-free water. The mix was incubated at 94°C for 1 min and the temperature was ramped down to 25°C (-0.1°C/s) in order to pre-anneal the oligos. For each reverse-transcriptase, a different reverse-transcription reaction was prepared with 50 ng poly(A) tailed RNA and 1 uL pre-annealed oligo. For Superscript II (Thermo, 18064014), RNA and oligos were mixed with 1uL dNTPs and RNase-free water up to 11 uL. Mixture was heated to 65°C for 5 mins and then quickly chilled on ice. Then 4 uL 5X First-strand buffer, 1 uL 0.1 M DTT, 1 uL RNAse Inhibitor Murine, and 1 uL SuperScript II were added to the mix. Reaction was then incubated at 42°C for 60 mins and inactivated by heating at 70°C for 15 mins. For Protoscript II (NEB, M0368S), the same reaction was performed with the 5X ProtoScript II buffer. For Superscript III (Thermo, 18080093), RNA and oligos were mixed with 1uL dNTPs, 4 uL 5X First-strand buffer, 1 uL 0.1 M DTT, 1 uL RNAse Inhibitor Murine, 1 uL Superscript III and RNase-free water up to 11 uL. Reaction was then incubated at 55°C for 60 mins and inactivated by heating at 70°C for 15 mins. For Superscript IV (Thermo, 18090010), RNA and oligos were mixed with 1uL dNTPs, 4 uL 5X SSIV buffer, 1 uL 0.1 M DTT, 1 uL RNAse Inhibitor Murine, 1 uL Superscript IV and RNase-free water up to 11 uL. Reaction was then incubated at 60°C for 30 mins and inactivated by heating at 80°C for 10 mins. For Maxima H Minus RT (Thermo, EP0752), RNA and oligos were mixed with 1 uL dNTPs, 4 uL 5X RT buffer, 1 uL RNAse Inhibitor Murine, 1 uL Maxima enzyme and RNase-free water up to 14 uL. Reaction was then incubated at 60°C for 30 mins and inactivated by heating at 85°C for 5 mins. For TGIRT (InGex), RNA and oligos were mixed with 4 uL 5X TGIRT Reaction Buffer (2.25 M NaCl, 25 mM MgCl2, 100 mM Tris-HCl, pH 7.5), 1 uL 0.1 M DTT, 1 uL RNAse Inhibitor Murine, 1 uL TGIRT and RNase-free water up to 9.75 uL. Mix was pre-incubated at room temperature for 30 minutes, then 1.25 uL dNTP was added. Reaction was then incubated at 60°C for 60 mins and inactivated by heating at 75°C for 15 mins. For Induro RT (NEB, M0681S), RNA and oligos were mixed with 1uL dNTPs, 4 uL 5X Induro buffer, 1 uL RNAse Inhibitor Murine, 1 uL Induro and RNase-free water up to 11 uL. Reaction was then incubated at 60°C for 30 mins and inactivated by heating at 95°C for 10 mins. For Marathon RT (Kerafast), RNA and oligos were mixed with 1uL dNTPs, 10 uL 2X Marathon buffer (400 mM KCl, 4 mM MgCl2, 100 mM Tris-HCl, pH 8.3, 10 mM DTT, 40% Glycerol), 1 uL RNAse Inhibitor Murine, 1 uL Marathon and RNase-free water up to 6 uL. Reaction was then incubated at 42°C for 30 mins and inactivated by heating at 95°C for 1 mins. For the manganese buffer conditions of Protoscript II, Superscript II, Superscript III, Superscript IV, Maxima, and TGIRT, buffers disclosed by the manufacturer were prepared by replacing Magnesium with the same amount of Manganese Chloride (Thermo, AAJ63150AD). For Induro, since the manufacturer does not disclose the RT buffer composition, the TGIRT buffer (with Manganese) was used. For Marathon RT, as suggested in literature [9], 2X Marathon RT buffer was prepared, replacing Magnesium with 1 mM Manganese. Once the reverse transcription reaction was performed, RNAse Cocktail was added to the mix in order to digest the RNA, and the mix was incubated at 37°C for 10 minutes. The reaction was then cleaned up using 0.8X AMPure XP Beads. In order to be able to ligate the sequencing adapters to the first cDNA strand, 1 µL 25 µM Comp_DNA was re-annealed to the 15 µL cDNA in a tube with 2.25 µL 0.1 M Tris pH 7.5, 2.25 µL 0.5 M NaCl and 2 µL nuclease-free water. The mix was incubated at 90°C for 1 minute and the temperature was ramped down to 25 °C (-0.1°C/s) in order to anneal the complementary to the first strand cDNA. The ligation of the barcode was performed by mixing 22.5 uL end-prepped DNA, 2.5 uL Native Barcode (NB), 25 uL Blunt/TA Ligation Master Mix (NEB, M0367S) and incubating the mix at room temperature for 10 mins. The reaction was then cleaned up using 0.8X AMPure XP Beads. Once all the barcoded libraries were pooled after elution, the ligation of the sequencing Adapter Mix (AMX) was performed by mixing the 65 uL pooled barcoded DNAs (up to 200 fmol), 5 uL AMX, 20 uL 5X NEBNext Quick Ligation Reaction Buffer and 10 uL Quick T4 DNA Ligase, and incubating the mix at room temperature for 10 mins. The reaction was cleaned up using 0.8X AMPure XP beads, using WSB Buffer for washing. The sample was then eluted in Elution Buffer (EB) and mixed with Sequencing Buffer (SQB) and Loading Beads (LB) prior to loading onto a primed R9.4.1 flowcell. The library was run on MinION flowcell with MinKNOW acquisition software version v.3.5.5. A detailed step-by-step DMS-FIRST-seq protocol is provided as an additional supplementary file, both for R9 and R10 flowcells (**File S1**). In this work, DMS-FIRST-seq libraries of *in vitro* DMS-modified RNAs were prepared using TGIRT in manganese buffer, whereas libraries of *in vivo* DMS-modified RNAs, Induro in manganese buffer was used (TGIRT is not commercially available any more).

### Analysis of dRNA-seq datasets

For the yeast ribosomal RNA data, reads were downloaded from European Nucleotide Archive (ENA) under accession number PRJEB37798. Fast5 reads were basecalled using Guppy v6, under the RNA basecalling model, using default parameters. Prior to mapping, uridines were converted to thymidines using the command "awk ’{ if (NR%4==2) { gsub(/U/,"T",$1); print $1 } else print }’ file.fastq". Reads were mapped using minimap2[58] version 2.26-r1175, with the default parameters to the *S. cerevisiae* 5s, 5.8s, 18s and 28s rRNA reference sequences.

For the yeast Nano-tRNAseq data, reads were downloaded from European Nucleotide Archive (ENA) under accession number PRJEB55684. Fast5 reads were basecalled using Guppy v6, under the RNA basecalling model, using default parameters. Prior to mapping, uridines were converted to thymidines using the command "awk ’{ if (NR%4==2) { gsub(/U/,"T",$1); print $1 } else print }’ file.fastq". Reads were mapped using BWA version 0.7.17-r1188, using custom parameters (bwa mem -W13 -k6 -xont2d -T20) to the *S. cerevisiae* tRNA reference sequences.

For the *in vitro* DMS treated samples, fast5 reads were basecalled using Guppy v6, under the RNA basecalling model, using default parameters. Prior to mapping, uridines were converted to thymidines using the command "awk ’{if (NR%4==2) {gsub(/U/,"T",$1); print $1 } else print }’ file.fastq". Reads were mapped using graphmap version v0.5.2 , with the default parameters to the in vitro RNA sequences (Tetrahymena ribozyme and lysine riboswitch RNAs).

All the mapped sam files were then converted to bam, sorted and indexed using samtools version 1.19. Mapping features such as base quality, mismatches, insertions and deletions were extracted for each position using in-house *bamtostats.sh* script. All reference files used in this work are available in the GitHub repository (https://github.com/novoalab/firstseq). Statistics for each sequencing run can be found on **Table S14**.

### Annotations

The list of tRNA modifications present in *S. cerevisiae* tRNAs was obtained from MODOMICS (https://iimcb.genesilico.pl/modomics/sequences/) and was retrieved on 21 September 2021. tRNA expression estimates from ONT-based *S. cerevisiae* tRNA sequencing were obtained from Nano-tRNAseq [59], and Illumina-based *S. cerevisiae* tRNA sequencing were obtained from mim-tRNAseq [2] (GSE152621).

### Analysis of dcDNA-seq datasets and FIRST-seq

For the dcDNA-seq and FIRST-seq data of the yeast total RNA runs from various reverse-transcriptases, fast5 reads were basecalled using Guppy v6, under the DNA high-accuracy (hac) basecalling model.

Then, for dcDNA-seq dataset, reads were mapped to two reference sequences using a two-step approach. Initially, reads were mapped to the long reference sequence, which includes non-tRNA references (*S. cerevisiae* non-coding RNA biotypes and in vitro transcripts) using minimap2 version 2.26-r1175, with the parameters -ax map-ont --MD -t 2. Mapped reads were filtered using samtools version 1.19 (view -Sb -bq 59), sorted, and indexed. Unmapped reads were extracted, and the remaining reads were filtered using seqkit [60]. The filtered reads were subsequently mapped to the short reference sequence, which includes *S. cerevisiae* tRNA references, using BWA version 0.7.17-r1188 (mem -W 13 -k 6 -x ont2d). Mapped reads were sorted and indexed using samtools. Both long and short mapped datasets were merged using Picard’s MergeSamFiles tool [61], with MSD=true and ASSUME_SORTED=false parameters. The final merged BAM files were indexed using samtools.

For FIRST-seq datasets, the same pipeline was performed, with an addition of filtering out for the reverse strand reads with the option of -f 0x10 in samtools view command.

To analyze reverse transcription (RT) stops, read start and end positions were extracted from the BAM files using the in-house read_ends.sh script. Mapping features such as base quality, mismatches, insertions and deletions were extracted for each position using in-house *bamtostats.sh* script. All reference files used in this work are available in the GitHub repository (https://github.com/novoalab/firstseq). Statistics for each sequencing run can be found on **Table S14**.

### Analysis of DMS-FIRST-seq datasets

Fast5 reads of DMS-FIRST-seq experiments were basecalled using Guppy v6, under the DNA high-accuracy (hac) basecalling model. For the DMS-FIRST-seq data of the *in vitro* RNAs, reads were mapped to the *in vitro* RNA sequences (*Tetrahymena* ribozyme and lysine riboswitch RNAs), using minimap2 version 2.26-r1175, with the parameters -ax map-ont --MD -t 2. All the mapped sam files were then converted to bam, sorted and indexed using samtools version 1.19. Mapping features such as base quality, mismatches, insertions and deletions were extracted for each position using in-house *bamtostats.sh* script. For the DMS-FIRST-seq data of the *in vivo E.coli* and MDA-MB-231 samples, reads were mapped to selected structured RNA references for each organism, using minimap2 version 2.26-r1175, with the parameters -ax map-ont --MD -t 2. All the mapped sam files were then converted to bam, sorted and indexed using samtools version 1.19. Mapping features such as base quality, mismatches, insertions and deletions were extracted for each position using in-house *bamtostats.sh* script. All reference files used in this work are available in the GitHub repository (https://github.com/novoalab/firstseq). Statistics for each sequencing run can be found on **Table S14.**

### LC-MS/MS sample preparation and data analysis

Samples were digested with 1 μl of the Nucleoside Digestion Mix (New England BioLabs, M0649S) and the mixture was incubated at 37 °C for 1 h. Samples (20ng) were analyzed using an Orbitrap Eclipse Tribrid mass spectrometer (Thermo Fisher Scientific, San Jose, CA, USA) coupled to an EASY-nLC 1200 (Thermo Fisher Scientific (Proxeon), Odense, Denmark). Ribonucleosides were loaded directly onto the analytical column and were separated by reversed-phase chromatography using a 50-cm column with an inner diameter of 75 μm, packed with 2 μm C18 particles (Thermo Fisher Scientific, ES903). Chromatographic gradients started at 93% buffer A and 3% buffer B with a flow rate of 250 nl/min for 5 minutes and gradually increased to 30% buffer B and 70% buffer A in 20 min. After each analysis, the column was washed for 10min with 0% buffer A and 100% buffer B. Buffer A: 0.1% formic acid in water. Buffer B: 0.1% formic acid in 80% acetonitrile. The mass spectrometer was operated in positive ionization mode with nanospray voltage set at 2.4 kV and source temperature at 305°C. For Parallel Reaction Monitoring (PRM) method the quadrupole isolation window was set to 0.7 m/z, and MSMS scans were collected over a mass range of m/z 50-450, with detection in the Orbitrap at resolution of 120,000. MSMS fragmentation of defined masses and schedule retention time **(Table S15**) was performed using HCD at NCE 20 (except stated differently, **Table S15),** the auto gain control (AGC) was set to “Standard” and a maximum injection time of 246 ms was used. In each PRM cycle, one full MS scan at resolution of 120,000 was acquired over a mass range of m/z 220-700 with detection in the Orbitrap mass analyzer. Auto gain control (AGC) was set to 1e5 and the maximum injection time was set to 50 ms. Serial dilutions were prepared using commercial pure ribonucleosides (0.005-150 pg, Carbosynth, Toronto Research Chemicals) in order to establish the linear range of quantification and the limit of detection of each compound ^[62]^ A mix of commercial ribonucleosides was injected before and after each batch of samples to assess instrument stability and to be used as external standard to calibrate the retention time of each ribonucleoside.

Acquired data were analyzed with the Skyline-daily software (v24.1.1.284) and extracted precursor areas of the ribonucleosides were used for quantification. The raw proteomics data have been deposited to the MetaboLights [63] repository with the dataset identifier REQ20250127208298.

### m¹A site prediction using a differential machine learning pipeline

To identify m¹A sites, we developed a differential machine learning pipeline that leverages the distinct error signatures induced by this modification in two different reverse transcriptases, TGIRT and SS3. We trained two separate Gradient Boosting (GB) classifiers on FIRST-seq data from S. cerevisiae total RNA. The positive training class consisted of features from known m¹A-modified rRNA and tRNA positions, while the negative class consisted of unmodified adenosine residues. Key features included per-base mismatch frequency and the relative frequencies of misincorporated T, C, and G nucleotides.

The dataset was split 80/20 for training and testing, stratified by modification status, and class imbalance was corrected using inverse weighting. Both models performed exceptionally well on the held-out test set (AUC = 1.000 for TGIRT; AUC = 0.981 for SS3). For prediction in human mitochondrial RNA, the trained classifiers were applied to their respective TGIRT and SS3 datasets to generate per-position m¹A probability scores. To minimize false positives, we implemented a stringent filtering pipeline. Potential single nucleotide polymorphisms (SNPs) and RNA editing sites were identified and removed by excluding positions with very high mismatch frequencies (>0.85) in both samples. A final m¹A Score was computed using a dynamic metric that prioritizes positions with either (i) concordantly high combined scores across both enzymes, or (ii) large differential scores between the two enzyme-specific predictions, thereby capturing the distinctive enzyme-dependent signature of m¹A modifications. All scripts used for model training, evaluation, and prediction are available in the public GitHub repository (https://github.com/novoalab/firstseq).

## Supporting information

Supplementary Fiile

## DATA AVAILABILITY

Data generated in this study, including basecalled FASTQ and processed data, has been deposited in GEO, under accession GSE288618.

## CODE AVAILABILITY

All code used in this manuscript has been deposited in GitHub (https://github.com/novoalab/firstseq).

## AUTHOR CONTRIBUTIONS

OB performed most wet lab experiments and bioinformatic analyses included in this work, and made all figures. GD contributed to validation experiments of DMS-probed *in vivo* RNA structures. JSM contributed with supervision of this work. IV and DI provided vivo DMS-probed samples from *E. coli* and human cell lines. EMN conceived and supervised the project. OB and EMN wrote the manuscript, with contributions from all authors.

## ACKNOWLEDGEMENTS

We would like to thank Prof. Martin A. Smith (Ramaciotti Center for Genomics, Australia) for his guidance and helpful discussions when starting this project, and for putting the fun into functional annotation. We would like to thank Prof. Jean Denis Beaudoin (UConn, USA) for providing the *Tetrahymena ribozyme* plasmid used in this work. We would like to thank all former and current Novoa lab members for their insightful discussions along these years. We are grateful to the CRG Core Technologies Programme for their support and assistance in this work.

## REFERENCES

1. Upton HE, Ferguson L, Temoche-Diaz MM, Liu X-M, Pimentel SC, Ingolia NT, et al. Low-bias ncRNA libraries using ordered two-template relay: Serial template jumping by a modified retroelement reverse transcriptase. Proc Natl Acad Sci U S A [Internet]. 2021;118:e2107900118. Available from: 10.1073/pnas.2107900118

2. Behrens A, Rodschinka G, Nedialkova DD. High-resolution quantitative profiling of tRNA abundance and modification status in eukaryotes by mim-tRNAseq. Mol Cell [Internet]. 2021;81:1802–15.e7. Available from: 10.1016/j.molcel.2021.01.028

3. Safra M, Sas-Chen A, Nir R, Winkler R, Nachshon A, Bar-Yaacov D, et al. The m1A landscape on cytosolic and mitochondrial mRNA at single-base resolution [Internet]. Nature. 2017. p. 251–5. Available from: 10.1038/nature24456

4. Xu H, Yao J, Wu DC, Lambowitz AM. Improved TGIRT-seq methods for comprehensive transcriptome profiling with decreased adapter dimer formation and bias correction. Sci Rep [Internet]. 2019;9:7953. Available from: 10.1038/s41598-019-44457-z

5. Nakano Y, Gamper H, McGuigan H, Maharjan S, Li J, Sun Z, et al. Genome-wide profiling of tRNA modifications by Induro-tRNAseq reveals coordinated changes. Nat Commun [Internet]. 2025;16:1047. Available from: 10.1038/s41467-025-56348-1

6. Scheepbouwer C, Aparicio-Puerta E, Gomez-Martin C, Verschueren H, van Eijndhoven M, Wedekind LE, et al. ALL-tRNAseq enables robust tRNA profiling in tissue samples. Genes Dev [Internet]. 2023;37:243–57. Available from: https://pmc.ncbi.nlm.nih.gov/articles/PMC10111867/

7. Qin Y, Yao J, Wu DC, Nottingham RM, Mohr S, Hunicke-Smith S, et al. High-throughput sequencing of human plasma RNA by using thermostable group II intron reverse transcriptases. RNA [Internet]. 2016;22:111–28. Available from: 10.1261/rna.054809.115

8. Mohr S, Ghanem E, Smith W, Sheeter D, Qin Y, King O, et al. Thermostable group II intron reverse transcriptase fusion proteins and their use in cDNA synthesis and next-generation RNA sequencing. RNA [Internet]. 2013;19:958–70. Available from: https://pubmed.ncbi.nlm.nih.gov/23697550/

9. Guo L-T, Adams RL, Wan H, Huston NC, Potapova O, Olson S, et al. Sequencing and structure probing of long RNAs using MarathonRT: A next-generation reverse transcriptase. J Mol Biol [Internet]. 2020;432:3338–52. Available from: 10.1016/j.jmb.2020.03.022

10. Xu H, Nottingham RM, Lambowitz AM. TGIRT-seq protocol for the comprehensive profiling of coding and non-coding RNA biotypes in cellular, extracellular vesicle, and plasma RNAs. Bio Protoc [Internet]. 2021;11:e4239. Available from: 10.21769/BioProtoc.4239

11. Gustafsson HT, Ferguson L, Galan C, Yu T, Upton H, Kaymak E, et al. Deep sequencing of yeast and mouse tRNAs and tRNA fragments using OTTR. Elife [Internet]. 2025;14. Available from: 10.7554/eLife.77616

12. Hauenschild R, Tserovski L, Schmid K, Thüring K, Winz M-L, Sharma S, et al. The reverse transcription signature of N-1-methyladenosine in RNA-Seq is sequence dependent. Nucleic Acids Res [Internet]. 2015;43:9950–64. Available from: 10.1093/nar/gkv895

13. Werner S, Schmidt L, Marchand V, Kemmer T, Falschlunger C, Sednev MV, et al. Machine learning of reverse transcription signatures of variegated polymerases allows mapping and discrimination of methylated purines in limited transcriptomes. Nucleic Acids Res [Internet]. 2020;48:3734–46. Available from: 10.1093/nar/gkaa113

14. Motorin Y, Muller S, Behm-Ansmant I, Branlant C. Identification of modified residues in RNAs by reverse transcription-based methods. Methods Enzymol [Internet]. 2007;425:21–53. Available from: 10.1016/S0076-6879(07)25002-5

15. Ryvkin P, Leung YY, Silverman IM, Childress M, Valladares O, Dragomir I, et al. HAMR: high-throughput annotation of modified ribonucleotides. RNA [Internet]. 2013;19:1684–92. Available from: 10.1261/rna.036806.112

16. 16. Novoa EM, Beaudoin J-D, Giraldez AJ, Mattick JS, Kellis M. Best practices for genome-wide RNA structure analysis: combination of mutational profiles and drop-off information [Internet]. Bioinformatics. bioRxiv; 2017. Available from: https://www.biorxiv.org/content/10.1101/176883v2

17. Zubradt M, Gupta P, Persad S, Lambowitz AM, Weissman JS, Rouskin S. DMS-MaPseq for genome-wide or targeted RNA structure probing in vivo. Nat Methods [Internet]. 2017;14:75–82. Available from: 10.1038/nmeth.4057

18. Siegfried NA, Busan S, Rice GM, Nelson JAE, Weeks KM. RNA motif discovery by SHAPE and mutational profiling (SHAPE-MaP). Nat Methods [Internet]. 2014;11:959–65. Available from: 10.1038/nmeth.3029

19. Yang M, Zhu P, Cheema J, Bloomer R, Mikulski P, Liu Q, et al. In vivo single-molecule analysis reveals COOLAIR RNA structural diversity. Nature [Internet]. 2022;609:394–9. Available from: 10.1038/s41586-022-05135-9

20. Aw JGA, Lim SW, Wang JX, Lambert FRP, Tan WT, Shen Y, et al. Determination of isoform-specific RNA structure with nanopore long reads. Nat Biotechnol [Internet]. 2020 [cited 2024 Feb 7];39:336–46. Available from: https://www.nature.com/articles/s41587-020-0712-z

21. Stephenson W, Razaghi R, Busan S, Weeks KM, Timp W, Smibert P. Direct detection of RNA modifications and structure using single-molecule nanopore sequencing. Cell genomics [Internet]. 2022 [cited 2024 Feb 7];2:100097. Available from: https://pubmed.ncbi.nlm.nih.gov/35252946/

22. Bohn P, Gribling-Burrer A-S, Ambi UB, Smyth RP. Nano-DMS-MaP allows isoform-specific RNA structure determination. Nat Methods [Internet]. 2023 [cited 2024 Feb 7];20:849–59. Available from: https://www.nature.com/articles/s41592-023-01862-7

23. Zhong J-Y, Niu L, Lin Z-B, Bai X, Chen Y, Luo F, et al. High-throughput Pore-C reveals the single-allele topology and cell type-specificity of 3D genome folding. Nat Commun [Internet]. 2023;14:1250. Available from: 10.1038/s41467-023-36899-x

24. Philpott M, Watson J, Thakurta A, Brown T Jr, Brown T Sr, Oppermann U, et al. Nanopore sequencing of single-cell transcriptomes with scCOLOR-seq. Nat Biotechnol [Internet]. 2021;39:1517–20. Available from: 10.1038/s41587-021-00965-w

25. Begik O, Diensthuber G, Liu H, Delgado-Tejedor A, Kontur C, Niazi AM, et al. Nano3P-seq: transcriptome-wide analysis of gene expression and tail dynamics using end-capture nanopore cDNA sequencing. Nat Methods [Internet]. 2023;20:75–85. Available from: 10.1038/s41592-022-01714-w

26. Singh M, Al-Eryani G, Carswell S, Ferguson JM, Blackburn J, Barton K, et al. High-throughput targeted long-read single cell sequencing reveals the clonal and transcriptional landscape of lymphocytes. Nat Commun [Internet]. 2019;10:3120. Available from: 10.1038/s41467-019-11049-4

27. Mitchell D, Cotter J, Saleem I, Mustoe AM. Mutation signature filtering enables high-fidelity RNA structure probing at all four nucleobases with DMS. Nucleic Acids Res [Internet]. 2023;51:8744–57. Available from: 10.1093/nar/gkad522

28. Rouskin S, Zubradt M, Washietl S, Kellis M, Weissman JS. Genome-wide probing of RNA structure reveals active unfolding of mRNA structures in vivo. Nature [Internet]. 2014;505:701–5. Available from: 10.1038/nature12894

29. Beaudoin J-D, Novoa EM, Vejnar CE, Yartseva V, Takacs CM, Kellis M, et al. Analyses of mRNA structure dynamics identify embryonic gene regulatory programs. Nat Struct Mol Biol [Internet]. 2018;25:677–86. Available from: 10.1038/s41594-018-0091-z

30. Leger A, Amaral PP, Pandolfini L, Capitanchik C, Capraro F, Miano V, et al. RNA modifications detection by comparative Nanopore direct RNA sequencing. Nat Commun [Internet]. 2021;12:7198. Available from: 10.1038/s41467-021-27393-3

31. Cruciani S, Delgado-Tejedor A, Pryszcz LP, Medina R, Llovera L, Novoa EM. De novo basecalling of RNA modifications at single molecule and nucleotide resolution. Genome Biol [Internet]. 2025;26:38. Available from: 10.1186/s13059-025-03498-6

32. Jenjaroenpun P, Wongsurawat T, Wadley TD, Wassenaar TM, Liu J, Dai Q, et al. Decoding the epitranscriptional landscape from native RNA sequences. Nucleic Acids Res [Internet]. 2021;49:e7. Available from: 10.1093/nar/gkaa620

33. Abebe JS, Price AM, Hayer KE, Mohr I, Weitzman MD, Wilson AC, et al. DRUMMER—rapid detection of RNA modifications through comparative nanopore sequencing. Bioinformatics [Internet]. 2022 [cited 2023 Jun 7];38:3113–5. Available from: https://academic.oup.com/bioinformatics/article-abstract/38/11/3113/6569078

34. Liu H, Begik O, Novoa EM. EpiNano: Detection of m6A RNA Modifications Using Oxford Nanopore Direct RNA Sequencing. Methods Mol Biol [Internet]. 2021;2298:31–52. Available from: 10.1007/978-1-0716-1374-0_3

35. Huang S, Zhang W, Katanski CD, Dersh D, Dai Q, Lolans K, et al. Interferon inducible pseudouridine modification in human mRNA by quantitative nanopore profiling. Genome Biol [Internet]. 2021;22:330. Available from: 10.1186/s13059-021-02557-y

36. Piechotta M, Naarmann-de Vries IS, Wang Q, Altmüller J, Dieterich C. RNA modification mapping with JACUSA2. Genome Biol [Internet]. 2022;23:115. Available from: 10.1186/s13059-022-02676-0

37. Smith AM, Jain M, Mulroney L, Garalde DR, Akeson M. Reading canonical and modified nucleobases in 16S ribosomal RNA using nanopore native RNA sequencing. PLoS One [Internet]. 2019;14:e0216709. Available from: 10.1371/journal.pone.0216709

38. Begik O, Lucas MC, Pryszcz LP, Ramirez JM, Medina R, Milenkovic I, et al. Quantitative profiling of pseudouridylation dynamics in native RNAs with nanopore sequencing. Nat Biotech [Internet]. 2021; Available from: 10.1038/s41587-021-00915-6

39. Liu H, Begik O, Lucas MC, Ramirez JM, Mason CE, Wiener D, et al. Accurate detection of m6A RNA modifications in native RNA sequences. Nat Commun [Internet]. 2019;10:4079. Available from: 10.1038/s41467-019-11713-9

40. Soneson C, Yao Y, Bratus-Neuenschwander A, Patrignani A, Robinson MD, Hussain S. A comprehensive examination of Nanopore native RNA sequencing for characterization of complex transcriptomes. Nat Commun [Internet]. 2019;10:3359. Available from: 10.1038/s41467-019-11272-z

41. Ura H, Togi S, Niida Y. Poly(A) capture full length cDNA sequencing improves the accuracy and detection ability of transcript quantification and alternative splicing events. Sci Rep [Internet]. 2022;12:10599. Available from: https://www.nature.com/articles/s41598-022-14902-7

42. Byrne A, Le D, Sereti K, Menon H, Vaidya S, Patel N, et al. Single-cell long-read targeted sequencing reveals transcriptional variation in ovarian cancer. Nat Commun [Internet]. 2024;15:6916. Available from: https://pmc.ncbi.nlm.nih.gov/articles/PMC11319652/

43. Liu-Wei W, van der Toorn W, Bohn P, Hölzer M, Smyth RP, von Kleist M. Sequencing accuracy and systematic errors of nanopore direct RNA sequencing. BMC Genomics [Internet]. 2024;25:528. Available from: 10.1186/s12864-024-10440-w

44. Schwartz S. m1A within cytoplasmic mRNAs at single nucleotide resolution: a reconciled transcriptome-wide map. RNA [Internet]. 2018;24:1427–36. Available from: 10.1261/rna.067348.118

45. Xiong X, Li X, Wang K, Yi C. Perspectives on topology of the human m1A methylome at single nucleotide resolution. RNA [Internet]. 2018;24:1437–42. Available from: 10.1261/rna.067694.118

46. Li X, Xiong X, Zhang M, Wang K, Chen Y, Zhou J, et al. Base-Resolution Mapping Reveals Distinct m1A Methylome in Nuclear- and Mitochondrial-Encoded Transcripts. Mol Cell [Internet]. 2017;68:993–1005.e9. Available from: 10.1016/j.molcel.2017.10.019

47. Dominissini D, Nachtergaele S, Moshitch-Moshkovitz S, Peer E, Kol N, Ben-Haim MS, et al. The dynamic N(1)-methyladenosine methylome in eukaryotic messenger RNA. Nature [Internet]. 2016;530:441–6. Available from: https://www.nature.com/articles/nature16998

48. Grozhik AV, Olarerin-George AO, Sindelar M, Li X, Gross SS, Jaffrey SR. Antibody cross-reactivity accounts for widespread appearance of m1A in 5’UTRs. Nat Commun [Internet]. 2019;10:5126. Available from: 10.1038/s41467-019-13146-w

49. Zhou H, Rauch S, Dai Q, Cui X, Zhang Z, Nachtergaele S, et al. Evolution of a reverse transcriptase to map N1-methyladenosine in human messenger RNA. Nat Methods [Internet]. 2019;16:1281–8. Available from: 10.1038/s41592-019-0550-4

50. Helm M, Motorin Y. Detecting RNA modifications in the epitranscriptome: predict and validate. Nat Rev Genet [Internet]. 2017;18:275–91. Available from: https://www.ncbi.nlm.nih.gov/pubmed/28216634

51. Kristen M, Plehn J, Marchand V, Friedland K, Motorin Y, Helm M, et al. Manganese ions individually alter the reverse transcription signature of modified ribonucleosides. Genes (Basel) [Internet]. 2020;11:950. Available from: 10.3390/genes11080950

52. Begik O, Mattick JS, Novoa EM. Exploring the epitranscriptome by native RNA sequencing. RNA [Internet]. 2022;28:1430–9. Available from: 10.1261/rna.079404.122

53. Zheng G, Qin Y, Clark WC, Dai Q, Yi C, He C, et al. Efficient and quantitative high-throughput tRNA sequencing. Nat Methods [Internet]. 2015;12:835–7. Available from: 10.1038/nmeth.3478

54. Shi J, Zhang Y, Tan D, Zhang X, Yan M, Zhang Y, et al. PANDORA-seq expands the repertoire of regulatory small RNAs by overcoming RNA modifications. Nat Cell Biol [Internet]. 2021;23:424–36. Available from: 10.1038/s41556-021-00652-7

55. Hardwick SA, Chen WY, Wong T, Deveson IW, Blackburn J, Andersen SB, et al. Spliced synthetic genes as internal controls in RNA sequencing experiments. Nat Methods [Internet]. 2016;13:792–8. Available from: 10.1038/nmeth.3958

56. Huang J, Wang G. Organelle-associated rRNA degradation. Bio Protoc [Internet]. 2019;9:e3255. Available from: 10.21769/BioProtoc.3255

57. Incarnato D, Morandi E, Simon LM, Oliviero S. RNA Framework: an all-in-one toolkit for the analysis of RNA structures and post-transcriptional modifications. Nucl Acids Res [Internet]. 2018;46:e97. Available from: 10.1093/nar/gky486

58. Li H. Minimap2: pairwise alignment for nucleotide sequences. Bioinformatics [Internet]. 2018;34:3094–100. Available from: 10.1093/bioinformatics/bty191

59. Lucas MC, Pryszcz LP, Medina R, Milenkovic I, Camacho N, Marchand V, et al. Quantitative analysis of tRNA abundance and modifications by nanopore RNA sequencing. Nat Biotech [Internet]. 2023 [cited 2024 Dec 19];42:72–86. Available from: https://www.nature.com/articles/s41587-023-01743-6

60. Shen W, Le S, Li Y, Hu F. SeqKit: A cross-platform and ultrafast toolkit for FASTA/Q file manipulation. PLoS One [Internet]. 2016;11:e0163962. Available from: 10.1371/journal.pone.0163962

61. Picard [Internet]. [cited 2025 Oct 29]. Available from: https://broadinstitute.github.io/picard/

62. Sugimoto N. Handbook of Chemical Biology of Nucleic Acids [Internet]. Springer Nature; 2023. Available from: https://play.google.com/store/books/details?id=7xrOEAAAQBAJ

63. Yurekten O, Payne T, Tejera N, Amaladoss FX, Martin C, Williams M, et al. MetaboLights: open data repository for metabolomics. Nucl Acids Res [Internet]. 2024;52:D640–6. Available from: 10.1093/nar/gkad1045

