## Supplementary Figures for "FIRST-seq: a nanopore-based cDNA sequencing platform for RNA modification and structure profiling"

**Figure S1. Comparison of dRNA-seq and dcDNA-seq "error" signatures on modified sites. (a)** Integrated Genomics Viewer (IGV) coverage track snapshots show dRNA-seq (upper tracks) and dcDNA-seq data at 12 different known modified positions from yeast rRNA and tRNA molecules. Colored boxes on top of the IGV tracks indicate the annotated modified sites. IGV tracks are coloured if the mismatch frequency of a given base is greater than 0.1, and shown as gray otherwise. **(b)** Boxplots comparing mismatch frequencies at different modification sites and their neighbouring sites across dRNA-seq and dcDNA-seq datasets. **(c)** Mismatch frequency comparisons for Watson-Crick-neutral modifications (left) and Watson-Crick-disrupting modifications (right) in dRNA-seq and dcDNA-seq.

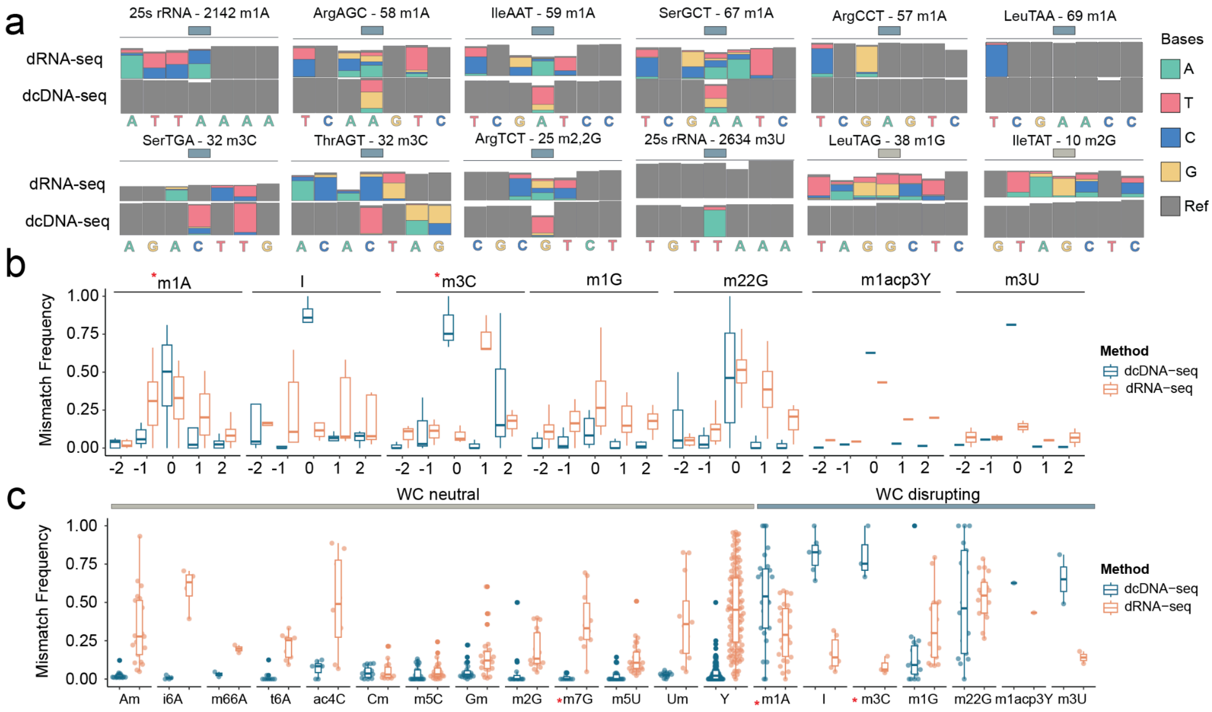

**Figure S2. Comparison of FIRST-seq and dcDNA-seq "error" signatures on modified sites. (a)** Density plot and line plot of (upper panels) 18s rRNA (lower panels) 25s rRNA, showing coverage and RT drop-off scores (based on read ends) for dcDNA-seq (left panels) and FIRST-seq (right panels). Watson-Crick disrupting modifications are labelled with a line under the coverage plot. We should note that sudden coverage drops observed in dcDNA-seq did not coincide with the modifications, but rather strand-switching artifacts, in agreement with previous works [1,2] **(b)** Barplot quantifying RT drop-off scores at specific modification sites across both methods. **(c)** Barplot showing aggregated "error" signatures from multiple Watson-Crick disrupting modification positions using dcDNA-seq (left panel) and FIRST-seq (right panel) data. **(d)** Scatter plot comparing mismatch frequencies between dcDNA-seq and FIRST-seq across Watson-Crick disrupting modification positions in tRNA and rRNAs. All experiments presented in this figure correspond to FIRST-seq and dcDNA-seq libraries performed on total yeast RNA. Panels a and b show a comparison of the two technologies on yeast rRNAs (18s and 25s). Panels c and d were generated using annotated yeast tRNA and rRNA modifications from the MODOMICS database [3]. To allow for direct comparison between FIRST-seq and dcDNA-seq, the FIRST-seq library was built using Maxima RT under $Mg^{2+}$ buffer conditions, matching the RT and buffer conditions used in the standard dcDNA-seq protocol.

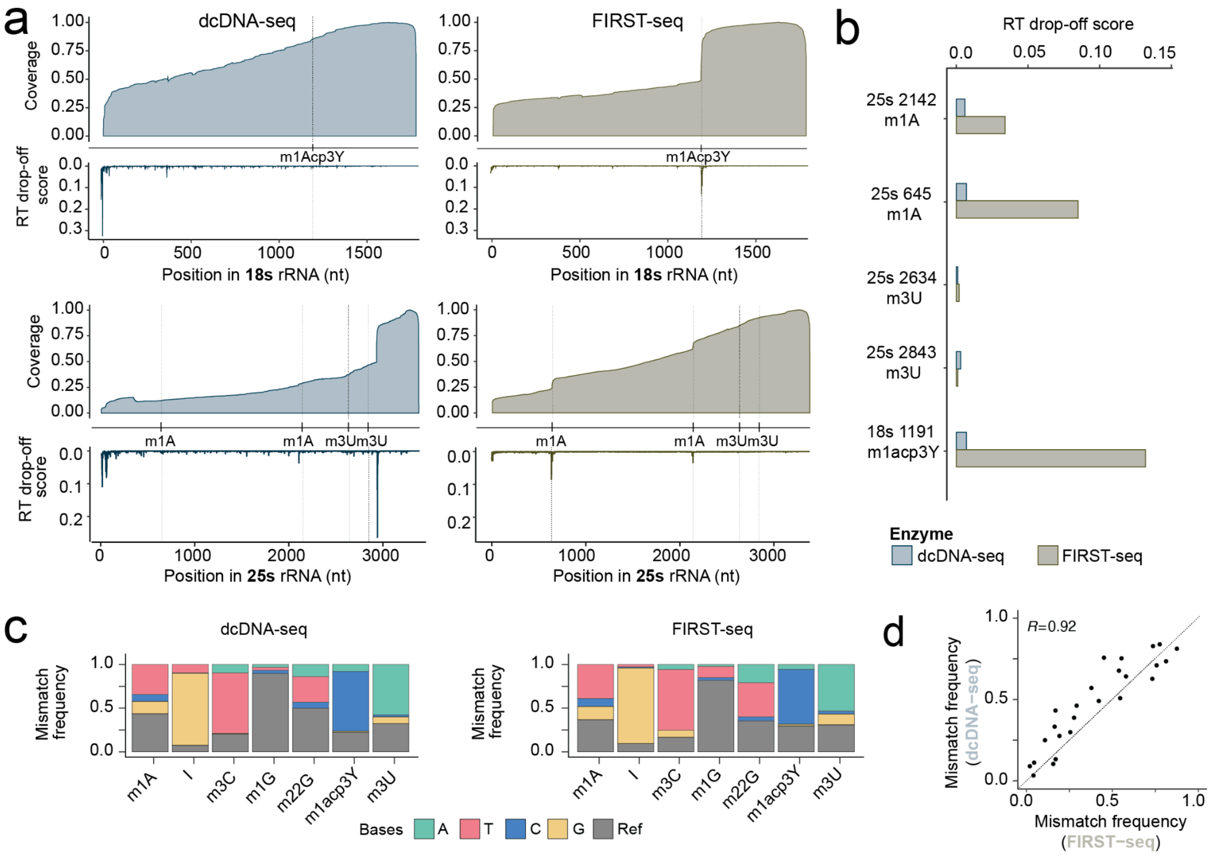

**Figure S3. FIRST-seq read lengths and read identity vary depending on RT enzyme and buffer of choice**. **(a)** TapeStation profiles of the RNA samples used in the study, including yeast total RNA, riboswitches, Tetrahymena ribozyme, and synthetic RNA sequences (Sequin controls). RNA template mix was then polyadenylated**. (b)** Electropherogram of fragment size distribution for RNA template mix. Peaks correspond to specific RNA fragment sizes, validating the quality of RNA samples used in subsequent experiments. **(c)** Integrated Genomics Viewer (IGV) coverage track snapshots show FIRST-seq data of the Yeast 25s rRNA from various reverse-transcriptases in magnesium ($MgCl_2$) and manganese ($MnCl_2$) buffers. Zoomed in boxes show the m$^1$A and m$^3$U modifications, which are known to cause RT drop-off. Mismatch frequency threshold for the coverage tracks is 0.1. **(d)** Barplot showing RT drop-off scores at specific 25S rRNA modification sites (m1A at positions 2142 and 645, m$^1$acp$^3$Y at position 1191, m$^3$U at positions 2634 and 2843) for each RT enzyme in $MgCl_2$ (top) and $MnCl_2$ (bottom) buffers. Colors match RT enzymes as indicated in the legend. **(e)** Barplot showing read length (nt) distribution of reads mapping to tRNAs (left panel) and ncRNA (right panel), obtained from FIRST-seq data of the yeast total RNA sequencing using various reverse-transcriptases in magnesium and manganese buffers.

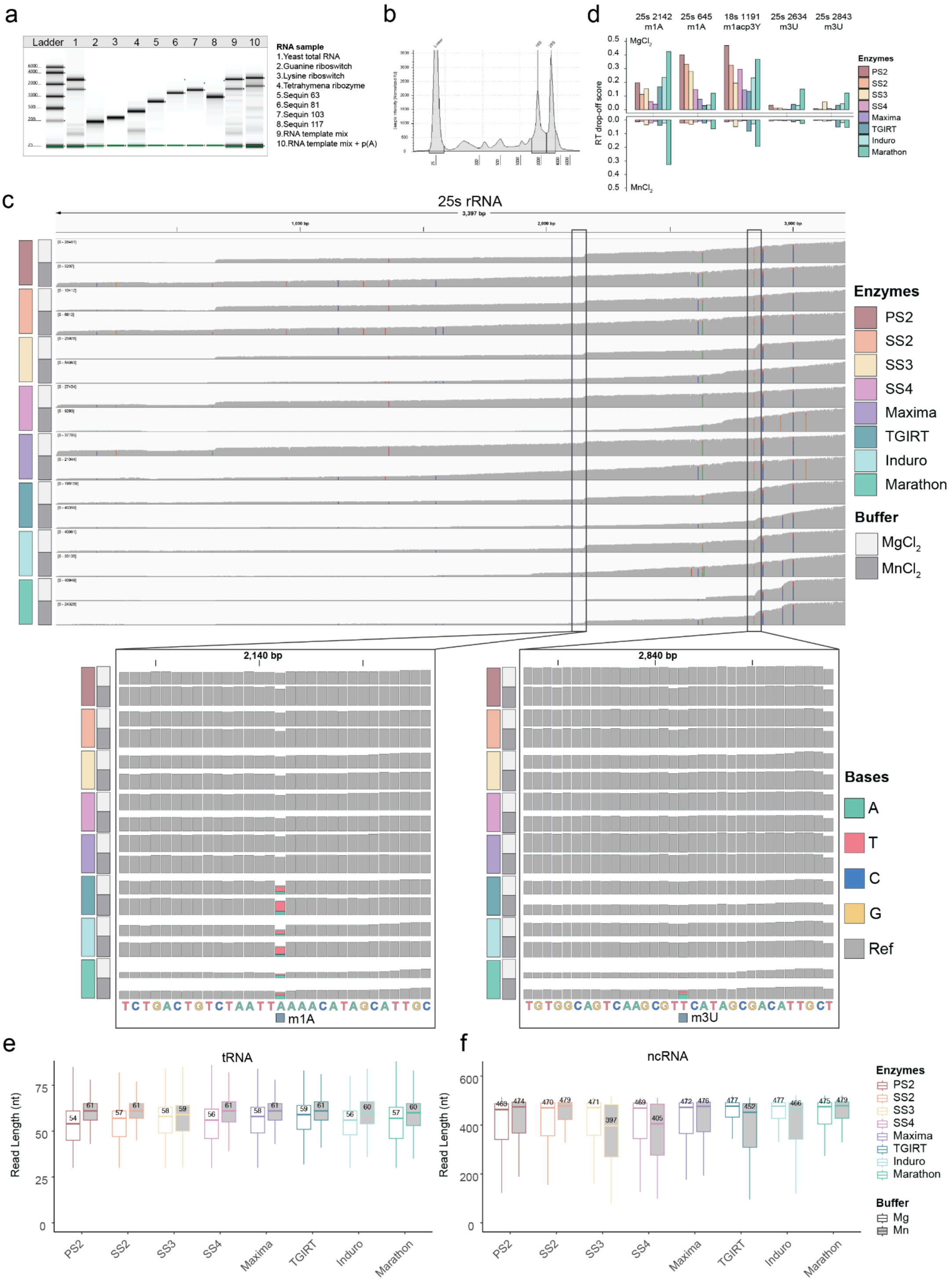

**Figure S4. Mismatch frequency analysis in *in vivo* DMS probing coupled to FIRST-seq in *E.coli* and *MDA-MB231* cell lines**. **(a)** Integrated Genomics Viewer (IGV) coverage track snapshots illustrate FIRST-seq data of unprobed and DMS-probed *E. coli* 16s ribosomal RNA sequenced using FIRST-seq. Zoomed box shows a specific region within the RNA, showing that higher DMS concentration correlates with increased mismatch errors at A and C positions. IGV allele frequency threshold for the coverage tracks is 0.05. **(b-c)** Boxplots showing overall mismatch frequency of DMS-FIRST-seq datasets of *E. coli* **(b)** and MDA-MB231 **(c)** DMS-probed total RNA, using different concentrations of DMS. Three biological replicates are shown. **(d-e)** Boxplots showing base-specific mismatch frequency of DMS-FIRST-seq datasets of the *E. coli* **(d)** and MDA-MB231 **(e)** total RNA, probed with different DMS concentrations. Three biological replicates shown. **(f)** Dotplot of AUC values calculated for the level of agreement between DMS reactivity scores obtained from FIRST-seq data using SS2 and Induro, and known RNA secondary structures for various RNAs in *E.coli* and *H.sapiens* (MDA-MB231) cells treated with 50 mM and 200 mM DMS. For the boxplots, the box extends from the first quartile to the third quartile of the data, with a line at the median. The whiskers extend from the box to the farthest data point lying within 1.5x the interquartile range from the box. Statistical analyses were performed using the Kruskal–Wallis test (p > 0.05:ns, p ≤ 0.05:*, p ≤ 0.01:**, p ≤ 0.001:***, p ≤ 0.0001:**** ).

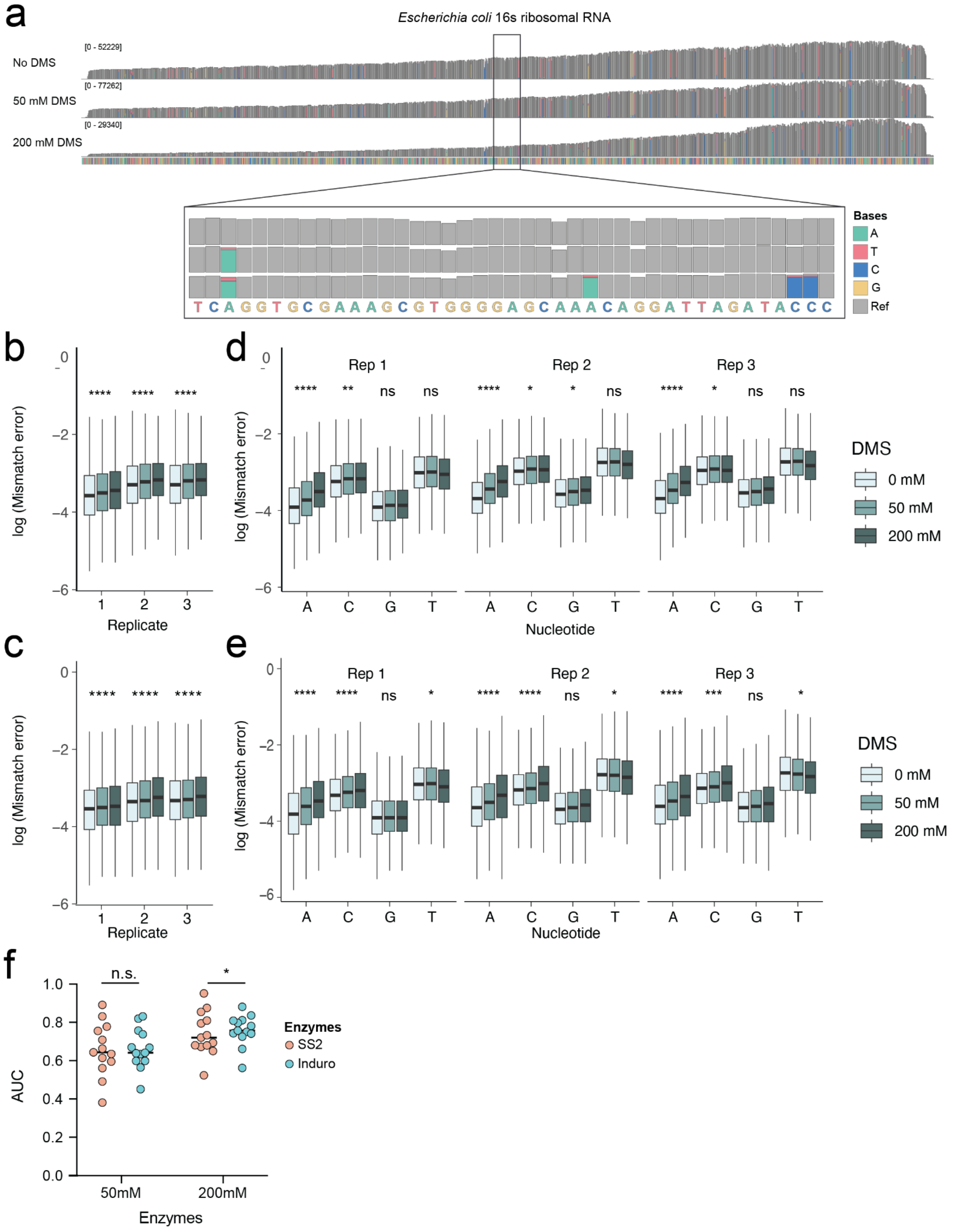

**Figure S****5****. Pairwise mismatch frequency comparison of enzymes and buffer conditions at rRNA and tRNA modification sites (a)** Scatter plot showing pairwise comparison of mismatch frequency of the enzymes in magnesium and manganese buffer conditions. Each dot represents a modified site in yeast rRNA and tRNA. **(b)** Scatter plot showing pairwise comparison of reverse transcriptases against SS3 in magnesium and manganese buffer conditions. Each dot represents a modified site in yeast rRNA and tRNA. Only the significantly changing modification types are colored for each scatter plot. Plots were generated using **Table S5.**

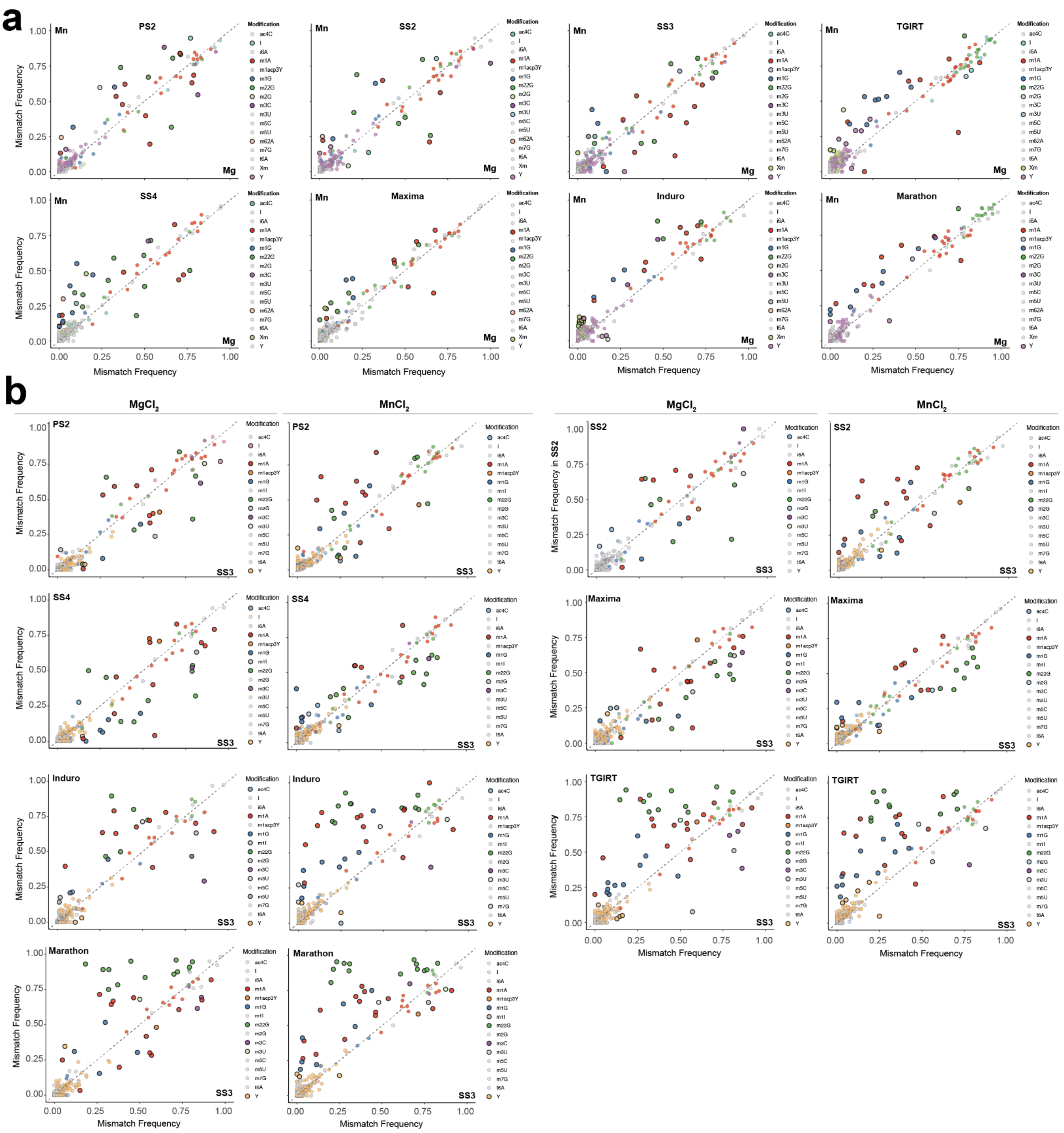

**Figure S6. PCA analysis of divergent error signatures of diverse reverse transcriptase enzymes in $MgCl_2$ and $MnCl_2$ buffers.** PCA plots showing the first two principal components (PC1 and PC2) based on mismatch signatures generated by different RT enzymes in $MgCl_2$ (left) and $MnCl_2$ (right) buffers. Each point represents an RT enzyme. The sites were taken from annotated RNA modifications in tRNAs and rRNAs used in the benchmarking RNA mix sample. Error signatures incorporate both mismatch frequency and relative base mismatch error composition.

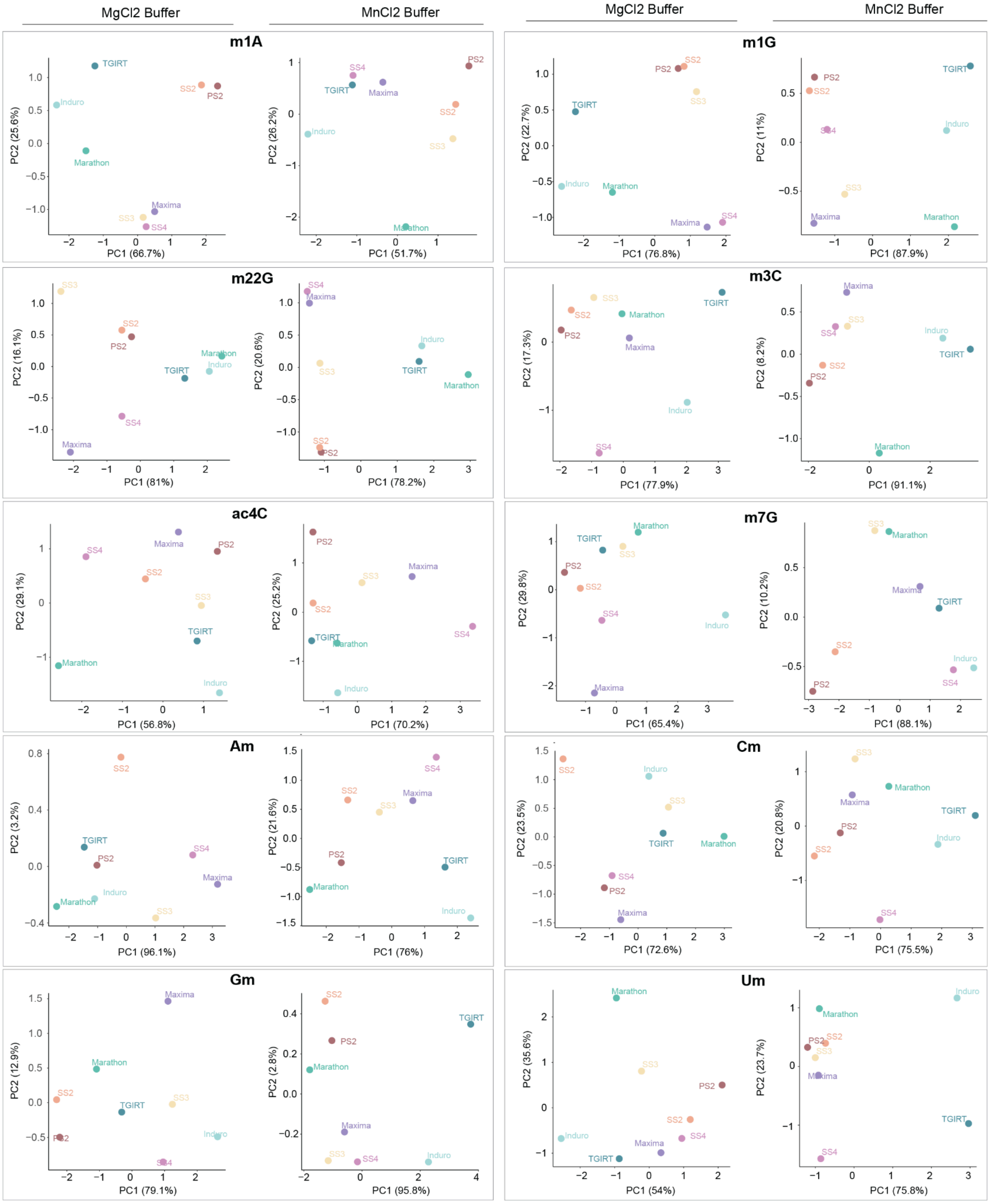

**Figure S7. Mismatch signatures of various RNA modifications and validation of the m¹A detection pipeline on mitochondrial ND5 mRNA.** Application of the m¹A detection pipeline to the human mitochondrial ND5 mRNA, which contains a known m¹A modification at position 1374. The top panels show Integrated Genomics Viewer (IGV) snapshots for TGIRT and SS3 libraries. The subsequent plots show the individual m¹A probability scores, RT drop-off scores, and combined scores for each enzyme. The final differential score (TGIRT-SS3 dynamic comparison) reveals a single, prominent peak precisely at the known m¹A site at position 1374.

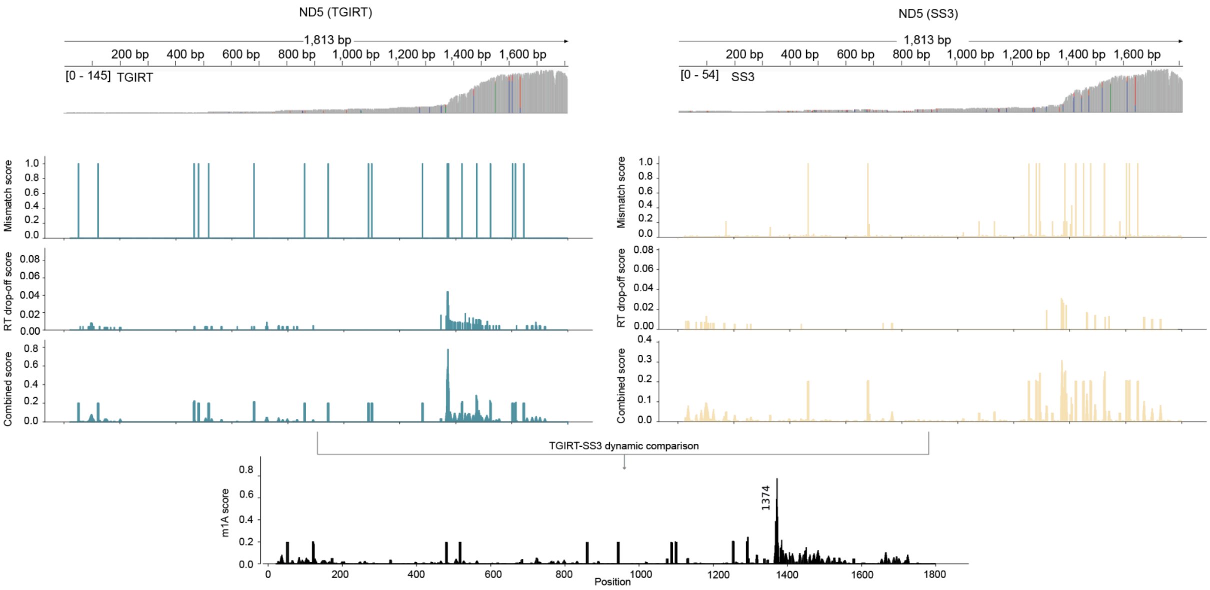

**Figure S8. RNA modifications generate distinct mismatch signatures depending on the RT enzyme and buffer choices.** Mismatch frequency profiles at various known modified sites in yeast tRNAs, including N³-methylcytidine (m³C), N¹-methylguanosine (m¹G), N²,N²-dimethylguanosine (m²²G), and inosine (I). The performance of eight different reverse transcriptases is compared under both magnesium and manganese buffer conditions. The color of the bars represents the base called at that position (A, T, C, G) or the reference base (Ref).

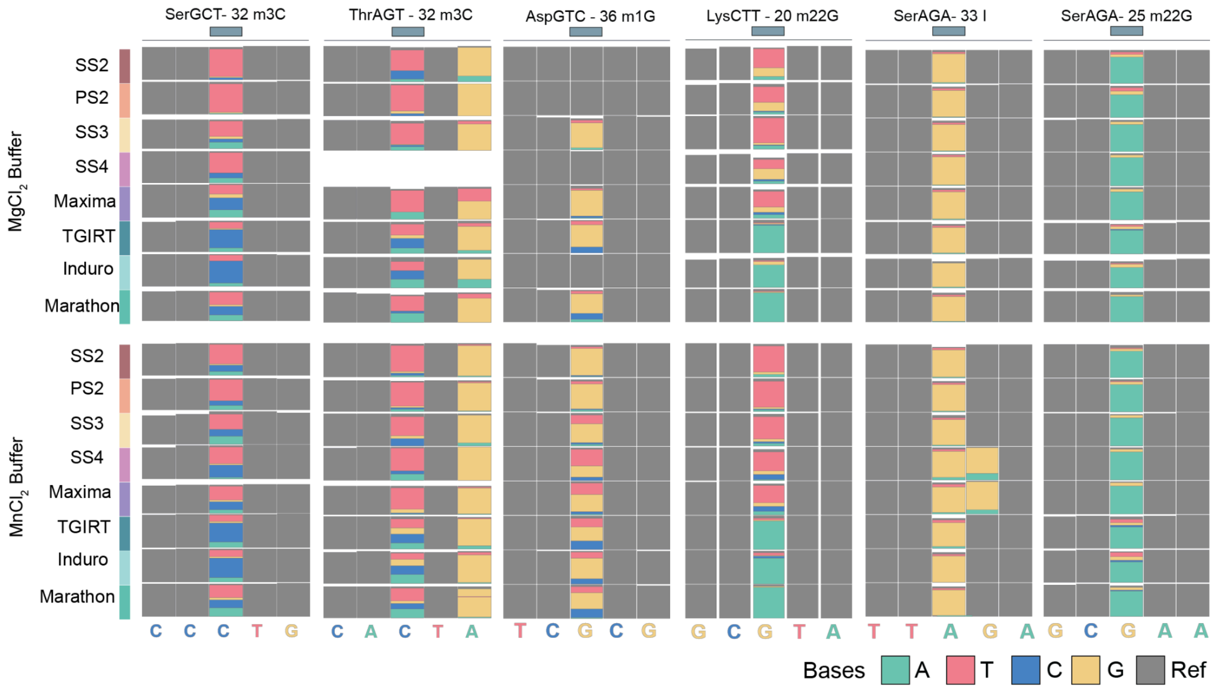

**Figure S9. Electronic gel images and mismatch frequency plots of *In vivo* data from *E.coli* and *MDA-MB231* cell lines. (a-b)** TapeStation profiles of the *E.coli* (a*)* and *MDA-MB231* cell line **(b)** RNA samples treated with different concentrations of DMS (0, 50, 200 mM, 500 mM) (left panels). Boxplots showing base-specific mismatch frequency if the *E.coli* **(a)** and *MDA-MB231* cell line **(b)** RNA samples were treated with different concentrations of DMS (0, 50, 200 mM, 500 mM) (right panels).

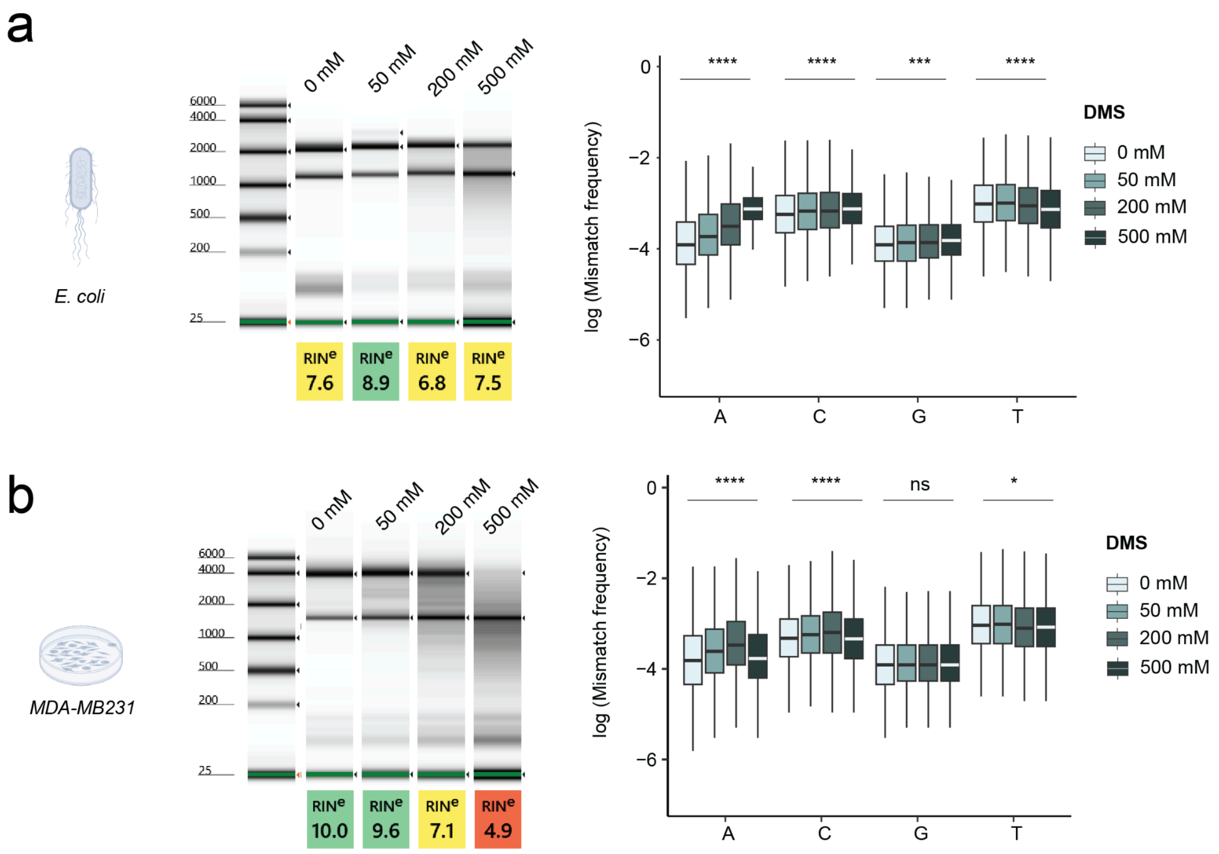

### SUPPLEMENTARY REFERENCES

1. Balázs Z, Tombácz D, Csabai Z, Moldován N, Snyder M, Boldogkői Z. Template-switching artifacts resemble alternative polyadenylation. BMC Genomics [Internet]. 2019 [cited 2024 Feb 12];20. Available from: https://www.ncbi.nlm.nih.gov/pmc/articles/PMC6839120/

2. Kainth AS, Haddad GA, Hall JM, Ruthenburg AJ. Merging short and stranded long reads improves transcript assembly. PLoS Comput Biol. 2023;19:e1011576.

3. Cappannini A, Ray A, Purta E, Mukherjee S, Boccaletto P, Moafinejad SN, et al. MODOMICS: a database of RNA modifications and related information. 2023 update. Nucleic Acids Res. 2024;52:D239–44.
