## Supplementary Fiile for "FIRST-seq: a nanopore-based cDNA sequencing platform for RNA modification and structure profiling"

### <u>SUPPLEMENTARY FILE S1: CONTENTS</u>


1. **FIRST-seq Protocol (R9 Chemistry)**
2. **FIRST-seq Protocol (R10 Chemistry)**

### 1. FIRST-seq Protocol (R9 Chemistry)

Notes before starting:

- The default protocol requires a polyA tail. In case you want to sequence RNAs that does not have polyA tail, you either need to PolyA tail your RNAs or use a sequence-specific oligo
- Volume for bead cleanup depends on the SIZE of your library. As a simple rule, more bead volume keeps shorter RNAs, but also keeps more adapters. Less bead volume removes enriches for longer RNAs, but if it's too little volume, might end up losing the RNA of interest too.
- This protocol assumes that the average length of the RNAs are around 2000 nts
- This protocol uses Direct cDNA-sequencing kit (DCS109) and Native Barcoding Expansion kit (EXP-NBD104) provided by Oxford Nanopore Technologies (ONT).

#### 1.1. Preannealing of the VNP and Complementary oligo

VNP: /5Phos/ACTTGCCTGTCGCTCTATCTTCTTTTTTTTTTTTTTTTTTTTVN
Comp_DNA : GAAGATAGAGCGACAGGCAAGTA

| Components | Initial Concentration | Desired Concentration | Volume |
|---|---|---|---|
| VNP | 10 uM | 2 uM | 2 ul |
| Comp_DNA | 10 uM | 2 uM | 2 ul |
| Tris pH 7.5 | 0.1 M | 0.01 M | 1 ul |
| NaCL | 0.5 M | 0.05 M | 1 ul |
| Water | | | 4 ul |
| Total | | | 10 ul |

- Heat the mixture for 94°C for 1 mins and ramp down to RT at 0.1°C/s (in PCR machine).

#### 1.2. Reverse Transcription

Note: This protocol includes the reaction for Induro enzyme, which is finally used in DMS-FIRST-seq. Users can replace this step with any reverse transcriptase reaction.

| Components | Volume |
|---|---|
| Preannealed oligos | 1 ul |
| RNA (100 ng) | Up to 12 ul |
| dNTPs (10 mM) | 1 ul |
| 5X Induro Buffer | 4 ul |
| RNase Inhibitor Murine | 1 ul |
| Induro RT | 1 ul |
| Total | 20 ul |

Incubate at **60°C** for **30 mins.**

Inactivate the reaction by heating at 95°C for 1minutes

##### 1.3. RNA Digestion and cleanup

- Add 1 ul RNAse Cocktail to each tube to digest the RNA strand
- Incubate at **37°C** for **10 mins**
- Move reaction to ice
- Mix the samples with the appropriate volume of beads (**0.8 X keeps everything above 150bp, good for getting rid of adapters**)
- Mix the beads by flicking
- Incubate 10 minutes at room temperature
- Spin down the tube and place it on the magnet
- Remove the supernatant
- Add 70% freshly prepared ethanol to the tube (200 ul)
- Incubate for 30 seconds at room temperature
- Remove the ethanol completely by spinning down and placing it back on magnet.
- Repeat the washing step
- Air-dry the pellet for maximum 1 minute, do not let it dry out completely!
- Resuspend the beads in 16 ul water
- Incubate 5-10 minutes in RT
- Place the beads on magnet
- Remove the elute and proceed to next step with it
- Use 1 ul for Qubit and TapeStation

##### 1.4. Annealing of complementary DNA to VNP oligo

*We need to reanneal the complementary DNA in case it dissociated in heat inactivation step

| Components | Initial Concentration | Desired Concentration | Volume |
|---|---|---|---|
| cDNA | | | 15 ul |
| Tris pH 7.5 | 0.1 M | 0.01 M | 2.25 |
| NaCL | 0.5 M | 0.05 M | 2.25 |
| Comp_DNA | 10uM | 1 uM | 2.25 ul |
| Water | | | 2 ul |
| Total | | | 22.5 ul |

- Mix by flicking!
- Heat the cDNA-oligo mix up to 90°C , incubate 1 minute
- Ramp down to 25°C - 0.1°C/s
- Mix the following :
    - 22.5 ul cDNA-complement mix
    - 2.5 ul Native Barcode
    - 25 ul Blunt/TA Ligase Mix

- Mix by flicking!
- Spin down
- Incubate the reaction for 10 minutes at room temperature

##### 1.5. Ampure XP Beads Cleanup

- Add 50 ul resuspended AMPure XP beads (1X, depends on the size of your library) to the reaction and mix by flicking
- Incubate 10 minutes at room temperature
- Spin down the tube and place it on the magnet
- Remove the supernatant
- Add 70% freshly prepared ethanol to the tube (200 ul)
- Incubate for 30 seconds at room temperature
- Repeat the washing step
- Remove the ethanol completely by spinning down and placing back it on magnet
- Air-dry the pellet for maximum 1 minute, do not let it dry out completely!
- Resuspend the beads in 16 ul water
- Incubate  5-10 minutes  at room temperature
- Place the beads on magnet
- Remove the elute and proceed to next step with it
- Quantify the cDNA with QUBIT DNA HS Assay
- Pool the barcoded samples at the desired ratio to a final volume of 65 µl in a DNA LoBind 1.5ml Eppendorf tube. Aim for as high a concentration as possible which does not exceed

200 fmoles total. If the total volume is >65 µl, perform a 2.5x AMPure clean up and elute in 65 µl of nuclease free water.

- Thaw ABB Buffer (ABB), Elution Buffer (EB) and NEBNext Quick Ligation Reaction Buffer (5x) at RT, mix by vortexing, spin down and place on ice. Check the contents or each tube are clear of any precipitate. Take out AMII (Adapter Mix II) and place on ice
- Check the contents of each tube are clear of any precipitate and are thoroughly mixed before setting up the reaction.
- Check that there is no precipitate present (DTT in the Blunt/TA Master Mix, if used, can sometimes form a precipitate)
- Spin down briefly before accurately pipetting the contents into the reaction
- Taking the pooled and barcoded DNA, perform adapter ligation as follows, mixing by flicking the tube between each sequential addition.
    - 65 µl 200 fmol pooled barcoded sample
    - 5 µl Adapter Mix II (AMII)
    - 20 µl NEBNext Quick Ligation Reaction Buffer (5X)
    - 10 µl Quick T4 DNA Ligase

- Mix gently by flicking the tube, and spin down.
- Incubate the reaction for 10 minutes at RT.
- Prepare the AMPure XP beads for use; resuspend by vortexing.
- Add 65 µl of resuspended AMPure XP beads **(Normally its 50 ul , 0.5X,** depends on the size of your library) to the reaction and mix by pipetting.
- Incubate on a Hula mixer (rotator mixer) for 5 minutes at RT.
- Place on magnetic rack, allow beads to pellet and pipette off supernatant.
- Add 140 **µl** of the ABB buffer to the beads. Close the tube lid, and resuspend the beads by flicking the tube. Return the tube to the magnetic rack, allow beads to pellet and pipette off the supernatant.
- **Repeat the previous step.**
- Remove the tube from the magnetic rack and resuspend pellet in 13 µl of Elution Buffer (EB).
- Pellet beads on magnet until the eluate is clear and colourless.
- Remove and retain 13 µl of eluate into a clean 1.5 ml Eppendorf DNA LoBind tube.
- Quantify 1 µl of eluted sample using a Qubit fluorometer
- Mix the contents of LB (Loading beads) with a large volume of pipette to make it homogeneous
- In the new tube, prepare the following:
    - 37.5 µl Sequencing Buffer (SQB)
    - 25.5 µl Loading Beads (LB), mixed immediately before use
    - 12 µl DNA library
- Mix the prepared library gently by pipetting up and down just prior to loading.

##### 1.6. Priming the flow cell and loading the library

- Take out the flowcell(s) from the fridge
- Insert it to the MinION device
- QC the flowcell(s) to ensure they are above the warranty (800 pores for MinION flowcells).
- Open the priming port by sliding it down clockwise
- Adjust 1000ul pipette to 200 ul

- Put the tip inside the priming port
- While the tip is in the port, increase the volume of pipette from 200ul up to 220ul
- Remove the tip from the port once you see a yellow liquid
- Thaw Running Buffer (RRB), Flush Buffer (FB) and Flush Tether (FLT)
- Mix FLT by pipetting and return to ice
- Mix RRB and FB by vortexing and return to ice
- Mix one tube of Flush Buffer (FB) and 30 ul Flush Tether  (FLT)
- Take 800 ul FB+FLT mix and introduce it to priming port:
    - To avoid bubbles, when the tip is really close to entering the priming port, put the first drop outside and then insert the tip to the port.
    - Insert all the mix slowly to the port AND put the last drop also outside
- Incubate 5 minutes while priming port is still open
- Open the sample port
- Add 200 ul more FB+FLT mix the same way with 1000 ul pipette to the priming port and see the bubbles coming out of the sample port
- Now put only one drop of the library through the sample port and start the sequencing
    - On the flow cell
        - Close both priming and sample ports
        - Close the lid
- Once you are sure that the flowcellis fine, apply the rest of the library by opening both the priming and sample ports and loading the library through the sample port. Close the ports again.

#### 2. FIRST-seq Protocol (R10 Chemistry)

Notes before starting:

- The default protocol requires a polyA tail. In case you want to sequence RNAs that does not have polyA tail, you either need to PolyA tail your RNAs or use a sequence-specific oligo
- Volume for bead cleanup depends on the SIZE of your library. As a simple rule, more bead volume keeps shorter RNAs, but also keeps more adapters. Less bead volume removes enriches for longer RNAs, but if it's too little volume, might end up losing the RNA of interest too.
- This protocol assumes that the average length of the RNAs are around 2000 nts
- This protocol needs Native Barcoding Expansion kit (SQK-LSK114).

##### 2.1. Preannealing of the VNP and Complementary oligo

VNP: /5Phos/ACTTGCCTGTCGCTCTATCTTCTTTTTTTTTTTTTTTTTTTTVN
Comp_DNA : GAAGATAGAGCGACAGGCAAGTA

| Components | Initial Concentration | Desired Concentration | Volume |
|---|---|---|---|
| VNP | 10 uM | 2 uM | 2 ul |
| Comp_DNA | 10 uM | 2 uM | 2 ul |
| Tris pH 7.5 | 0.1 M | 0.01 M | 1 ul |
| NaCL | 0.5 M | 0.05 M | 1 ul |
| Water | | | 4 ul |
| Total | | | 10 ul |

- Heat the mixture for 94°C for 1 mins and ramp down to RT at 0.1°C/s (in PCR machine).

##### 2.2. Reverse Transcription

Note: This protocol includes the reaction for Induro enzyme, which is finally used in DMS-FIRST-seq. Users can replace this step with any reverse transcriptase reaction.

| Components | Volume |
|---|---|

| | |
|---|---|
| Preannealed oligos | 1 ul |
| RNA (100 ng) | Up to 12 ul |
| dNTPs (10 mM) | 1 ul |
| 5X Induro Buffer | 4 ul |
| RNase Inhibitor Murine | 1 ul |
| Induro RT | 1 ul |
| Total | 20 ul |

Incubate at **60°C** for **30 mins.**

Inactivate the reaction by heating at 95°C for 1minutes

##### 2.3. RNA Digestion and cleanup

- Add 1 ul RNAse Cocktail to each tube to digest the RNA strand
- Incubate at **37°C** for **10 mins**
- Move reaction to ice
- Mix the samples with the appropriate volume of beads (**0.8 X keeps everything above 150bp, good for getting rid of adapters**)
- Mix the beads by flicking
- Incubate 10 minutes at room temperature
- Spin down the tube and place it on the magnet
- Remove the supernatant
- Add 70% freshly prepared ethanol to the tube (200 ul)
- Incubate for 30 seconds at room temperature
- Remove the ethanol completely by spinning down and placing it back on magnet.
- Repeat the washing step
- Air-dry the pellet for maximum 1 minute, do not let it dry out completely!
- Resuspend the beads in 16 ul water
- Incubate 5-10 minutes in RT
- Place the beads on magnet
- Remove the elute and proceed to next step with it
- Use 1 ul for Qubit and TapeStation

##### 2.4. Annealing of complementary DNA to VNP oligo

*We need to reanneal the complementary DNA in case it dissociated in heat inactivation step

| **Components** | **Initial Concentration** | **Desired Concentration** | **Volume** |
|---|---|---|---|

| | | | |
|---|---|---|---|
| cDNA | | | 15 ul |
| Tris pH 7.5 | 0.1 M | 0.01 M | 2.25 |
| NaCL | 0.5 M | 0.05 M | 2.25 |
| Comp_DNA | 10uM | 1 uM | 2.25 ul |
| Water | | | 2 ul |
| Total | | | 22.5 ul |

- Mix by flicking!
- Heat the cDNA-oligo mix up to 90°C , incubate 1 minute
- Ramp down to 25°C - 0.1°C/s
- Mix the following :
    - 22.5 ul cDNA-complement mix
    - 2.5 ul Native Barcode (NB01-24, ONT)
    - 25 ul Blunt/TA Ligase Mix

- Mix by flicking!
- Mix by flicking!
- Incubate at room temperature for 20 minutes. In the meantime, equilibrate the Ampure XP beads at least 30 minutes before use.
- Add 5 ul of EDTA (0.5 M, pH 8) to each tube and mix thoroughly by pipetting and quickly spin down to inactive the reaction.
- Pool all the barcoded samples in a 1.5 mL DNA LoBind tube and proceed to cleanup

#### 2.5. Ampure XP Beads Cleanup

- Add 50 ul resuspended AMPure XP beads (1X, depends on the size of your library) to the reaction and mix by flicking
- Incubate 10 minutes at room temperature
- Spin down the tube and place it on the magnet
- Remove the supernatant
- Add 70% freshly prepared ethanol to the tube (200 ul)
- Incubate for 30 seconds at room temperature
- Repeat the washing step
- Remove the ethanol completely by spinning down and placing back it on magnet
- Air-dry the pellet for maximum 1 minute, do not let it dry out completely!
- Resuspend the beads in 31 ul water
- Incubate  5-10 minutes  at room temperature
- Place the beads on magnet
- Remove the elute and proceed to next step with it
- Quantify the cDNA with Qubit DNA HS Assay

#### 2.6. Native Adapter (NA) Ligation

In a 1.5 ml Eppendorf LoBind tube, mix in the following order:

| Reagent | Volume |
|---|---|
| Pooled barcoded sample (Up to 200 fmol) | 30 µl |
| Native Adapter (NA) | 5 µl |
| NEBNext Quick Ligation Reaction Buffer (5X) | 10 µl |
| Quick T4 DNA Ligase | 5 µl |
| **Total** | **50 µl** |

●Mix gently by flicking the tube, and spin down.
●Incubate the reaction for 20 minutes at RT.

Note: Depending on the wash buffer (LFB or SFB) used, the clean-up step after adapter ligation is designed to either enrich DNA fragments of >3 kb, or purify all fragments equally.
To enrich for DNA fragments of 3 kb or longer, use Long Fragment Buffer (LFB)
To retain DNA fragments of all sizes, use Short Fragment Buffer (SFB)

#### 2.7. Ampure XP Beads Cleanup

- Add 25 ul resuspended AMPure XP beads (0.5X, depending on the size of library) to the reaction and mix by flicking
- Incubate 10 minutes at room temperature
- Thaw Long or Short Fragment Buffer (LFB, SFB) and Elution Buffer (EB) at RT, mix by vortexing, spin down and place on ice. Check if the contents of each tube are clear of any precipitate.
- Spin down the tube and place it on the magnet
- Remove the supernatant
- Add 125 ul SFB or LFB to the beads. Close the tube lid, and resuspend the beads by flicking. Return the tube to the magnetic rack, allow beads to pellet and pipette off the supernatant
- Repeat the previous step.

- Remove the SFB/LFB completely by spinning down the tube, pipetting out the liquid and placing back it on magnet
- Air-dry the pellet for maximum 1 minute, do not let it dry out completely!
- Resuspend the beads in 15 ul Elution Buffer (EB)
- Incubate 5-10 minutes in RT
- Place the beads on magnet
- Place the elute in a 1.5 ml tube
- Quantify 1 µl of eluted sample using a Qubit fluorometer
- Make the library up to 12 µl at 10-20 fmol.
- We recommend loading 10 - 20 fmol of this final prepared library onto the R10 flow cell.
- The prepared library is used for loading into the MinION flow cell. Store the library on ice until ready to load.
- Move to the “Prepare the library for loading” step.

##### 2.8. Prepare the library for loading and prime the flow cell

Before starting:

- Thaw the Sequencing Buffer (SB), Library Beads (LIB) or Library Solution (LIS, if using), Flow Cell Tether (FCT) and one tube of Flow Cell Flush (FCF) at RT. Mix by vortexing, spin down and return to ice.
- For optimal sequencing performance and improved output on MinION R10.4.1 flow cells (FLO-MIN114), we recommend adding Bovine Serum Albumin (BSA) to the flow cel priming mix at a final concentration of 0.2 mg/ml
- Mix the contents of LIB (Library beads) with a large volume of pipette to make it homogeneous

- To prepare the flow cell priming mix with BSA, add the following directly to the tube of Flow cell Flush and mix by pipetting:

  1,170 µl FCF
  5 ul BSA at 50 mg/ml
  30 ul FCT

- In the new tube, prepare the following:

  37.5 µl Sequencing Buffer (B)
  25.5 µl Loading Beads (LB), mixed immediately before use
  12 µl DNA library

<u>QC the flowcell:</u>

- Take the flowcell out of the fridge.Connect it to the MinION
- Check Flowcell (QC it to see how many active pores there are)

- Keep it connected until primed.

Prime the flowcell:

- Remove the bubble from the port
- Open the lid of the nanopore sequencing device and slide the flow cell's priming port cover clockwise so that the priming port is visible.
- Set a P1000 pipette to 200 µl
- Insert the tip into the priming port
- Turn the wheel until the dial shows 220-230 µl, or until you can see a small volume of buffer entering the pipette tip
- Load 800 ul of flow cell priming mix
- Wait for 5 minutes
- Open sample port cover
- Load 200 ul of priming mix more into the priming port, observing the bubbles coming out of sample port

Load the library:

- Mix the prepared library gently by pipetting up and down just prior to loading.
- Add 75 µl of sample to the flow cell via the SpotON sample port in a dropwise fashion. Ensure each drop flows into the port before adding the next.
- Gently replace the SpotON sample port cover, making sure the bung enters the SpotON port, close the priming port and replace the MinION lid.
